# Molecular basis for shifted receptor recognition by an encephalitic arbovirus

**DOI:** 10.1101/2025.01.01.631009

**Authors:** Xiaoyi Fan, Wanyu Li, Jessica Oros, Jesse S. Plung, Jessica A. Plante, Himanish Basu, Sivapratha Nagappan-Chettiar, Joshua M. Boeckers, Laurentia V. Tjang, Colin J. Mann, Vesna Brusic, Tierra K. Buck, Haley Varnum, Pan Yang, Linzy M. Malcolm, So Yoen Choi, William M. de Souza, Isaac M. Chiu, Hisashi Umemori, Scott C. Weaver, Kenneth S. Plante, Jonathan Abraham

**Author notes:** These authors contributed equally (Xiaoyi Fan, Wanyu Li). Correspondence (J.A.).

## Abstract

After decades of inactivity throughout the Americas, western equine encephalitis virus (WEEV) recently re-emerged in South America, causing a large-scale outbreak in humans and horses. WEEV binds protocadherin 10 (PCDH10) as a receptor; however, nonpathogenic strains no longer bind human or equine PCDH10 but retain the ability to bind avian receptors. Highly virulent WEEV strains can also bind the very low-density lipoprotein receptor (VLDLR) and apolipoprotein E receptor 2 (ApoER2) as alternative receptors. Here, by determining cryo-electron microscopy structures of WEEV strains isolated from 1941–2005 bound to mammalian receptors, we identify polymorphisms in the WEEV spike protein that explain shifts in receptor dependencies and that can allow nonpathogenic strains to infect primary cortical neurons. We predict the receptor dependencies of additional strains and of a related North American alphavirus. Our findings have implications for outbreak preparedness and enhance understanding of arbovirus neurovirulence through virus receptor binding patterns.

## Introduction

Several alphaviruses cause outbreaks of encephalitis in humans and equids with unpredictable frequency and scale. These include western equine encephalitis virus (WEEV), eastern equine encephalitis virus (EEEV), and Venezuelan equine encephalitis virus (VEEV).^1–3^ Manifestations of WEEV infection range from mild or asymptomatic illness to encephalitis, and survivors can be left with devastating neurological sequalae that include seizures and cognitive impairment.^3^ WEEV previously caused large outbreaks of encephalitis in horses and humans in the early 20^th^ century in North America, but outbreaks have since drastically decreased in both frequency and scale (a phenomenon known as viral “submergence”).^4^ The last documented human case of WEEV in North America was in 1999, although the virus can still be detected in mosquito pools. Until very recently, the last outbreak of WEEV in South America was more than three decades ago,^5^ with only sporadic spillover events observed thereafter. For example, there was a single fatal human case in Uruguay in 2009, even though the presence of WEEV in Uruguay had not been documented since the 1970s.^6^ However, from November of 2023 to April of 2024, WEEV unexpectedly re-emerged in South America, causing a large-scale outbreak that involved over 2,500 equine cases, over 200 human cases, and resulted in a total of 12 human fatalities in Argentina, Uruguay, and Brazil.^7–9^

WEEV binds protocadherin 10 (PCDH10), a membrane protein that is broadly expressed but whose expression is particularly enriched in the brain, as a cellular receptor on mammalian cells.^10,11^ PCDH10 is a non-clustered δ2 protocadherin that is involved in synapse development.^12,13^ The PCDH10 ectodomain contains six extracellular cadherin repeats (EC1–6) that are connected by loops that are rigidified by calcium coordination (Figure 1A).^14^ The WEEV spike protein binds PCDH10 EC1, which is the most membrane-distal repeat in the receptor ectodomain.^10,11^ PCDH10 shares no structural homology with any previously described alphavirus receptors (e.g., matrix remodeling-associated protein 8 (MXRA8) or low-density lipoprotein receptor (LDLR)-related proteins);^15–22^ therefore, how PCDH10 interacts with alphavirus spike proteins is unknown.

**Figure 1.**
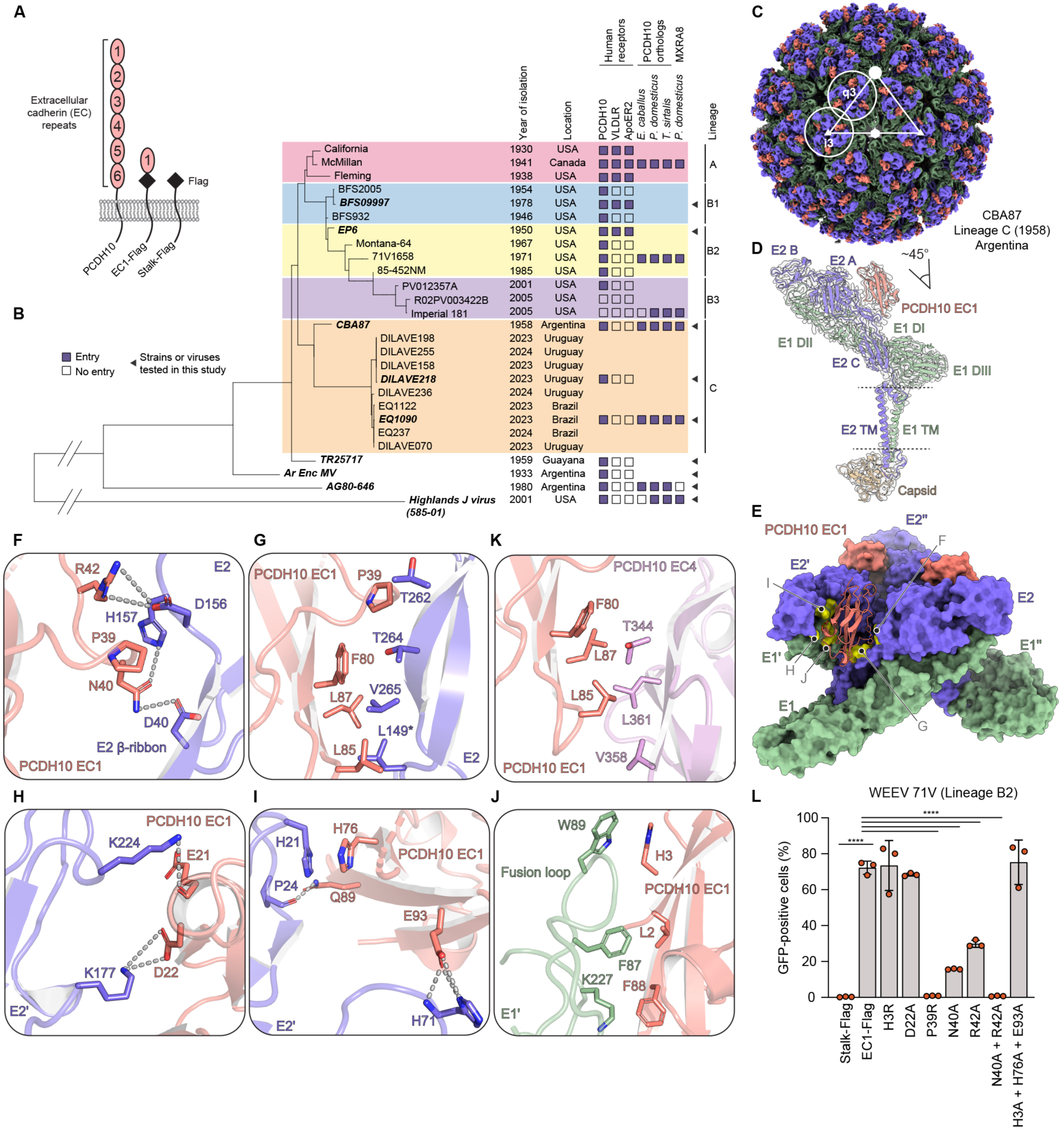
Structural basis for WEEV recognition of human PCDH10. (A) Schematic diagrams of PCDH10 and Flag-tagged constructs. (B) Partial phylogenetic tree and summary of infectivity assays with GFP-expressing RVPs for various WEEV strains and Highlands J virus in K562 cells stably expressing the indicated receptors. Strains newly tested in this study are indicated with black triangles. Other results were previously reported in Li et al.^10^ (C) Cryo-EM maps of human PCDH10_EC1_–Fc bound to WEEV CBA87 VLP. WEEV E2 and E1 and PCDH10 EC1 are shown in purple, green, and pink, respectively. The 5-fold (i5), 3-fold (i3), and 2-fold (i2) icosahedral symmetry axes are indicated with a closed circle, triangle, and hexagon, respectively. The icosahedral threefold (i3) and quasi-three-fold (q3) spikes are circled and indicated. (D) Ribbon diagram of a single WEEV E2–E1 protomer and PCDH10 EC1 repeat fitted into its associated cryo-EM density map. The angle with which EC1 is inserted into the cleft relative to the threefold axis of spike protein trimer is indicated. E2 and E1 domains are indicated. Dashed lines indicate the position of the viral membrane. TM: transmembrane. (E) Surface representation of the PCDH10 EC1-bound WEEV CBA87 spike protein. One of the three PCDH10 EC1 molecules is shown in ribbon representation. Clusters of spike protein residues that interact with EC1 are shown in yellow. Details for the indicated interacting regions are shown in panels F to J. (F–J) PCDH10 EC1 interactions with the WEEV E2–E1 protomer (F and G) and E2’–E1’ protomer (H-J). WEEV E2 L149, a key polymorphic residue, is indicated with an asterisk. Hydrogen bonds and salt bridges are shown as dashed lines. (K) Interaction interface of human PCDH10 EC1 and EC4 in a crystal structure of the PCDH10 EC1–EC4 homodimer (PDB: 6VFW).^14^ (L) Infection of K562 cells stably expressing wild-type and mutant human PCDH10 EC1 constructs by GFP-expressing WEEV RVP (strain 71V1658). Infection was quantified by flow cytometry. See Figure S8 for flow cytometry gating strategy and cell surface receptor staining. Data are mean ± s.d. from three experiments performed in duplicates or triplicates (n=3) (L). One-way ANOVA with Dunnett’s multiple comparisons test, \*\*\*\**P*<0.0001 compared to stalk-Flag (L).

WEEV strains isolated in North America are divided into lineages A and B; lineage B is further subdivided into B1, B2, and B3 (Figures 1B and S1; Table S1). Later lineages displaced earlier ones, and B3 is the most recently detected sublineage in North America.^4^ While human PCDH10 is a general receptor for most WEEV strains we previously tested, lineage A strains, which are highly virulent in animal models and were isolated in North America in the 1930s– 1940s, also bind VLDLR and ApoER2 (Figure 1B).^10,23^ Imperial 181 is a B3 WEEV strain that was isolated from a mosquito pool in California in 2005 and is nonpathogenic in animal models.^4,24^ Despite not binding to human or murine PCDH10,^10^ Imperial 181 can still bind PCDH10 orthologs of WEEV’s enzootic hosts, which include house sparrows (*Passer domesticus*). Imperial 181 can also bind the ortholog of common garter snakes (*Thamnophis sirtalis*), which are proposed overwintering hosts.^10,25,26^ The molecular basis for shifting patterns of receptor binding by different WEEV strains isolated in North America over the past century is unknown, nor are the receptor-binding properties of WEEV strains that recently re-emerged in South America.

Here, we conducted in depth structural and functional analyses of WEEV interactions with its alternative cellular receptors to identify the spike protein polymorphisms that explain the dynamic patterns of receptor recognition during the evolution of this important human pathogen.

## Results

### Structure of WEEV bound to human PCDH10

The alphavirus genome encodes four nonstructural proteins and structural proteins capsid, E3, E2, 6K, TF, and E1.^2^ The viral spike proteins E2 and E1 are arranged with icosahedral symmetry on the surface of virions.^27–29^ E2 and E1 form heterodimers that assemble as 80 trimers that bind receptors and mediate the fusion of viral and cellular membranes during viral entry.^27–29^ For structural analysis, we produced WEEV CBA87 virus-like particles (VLPs) (Figure S2A). This strain was previously isolated from the brain of an infected horse in Córdoba, Argentina, in 1958 and has been used in investigational VLP-based vaccines that were modified for high-yield expression.^30^ In biolayer interferometry experiments, monomeric soluble EC1 interacted with immobilized WEEV CBA87 VLPs with a K_D_ of 5.6 µM (Figures S2A–E). This affinity is similar to that of the PCDH10 ectodomain for itself during homodimerization (reported K_D_ of 3.6 µM).^14^

We determined cryo-EM structures of CBA87 VLPs alone or bound to PCDH10 EC1 (Figures 1C–E, S3A–F, and S4A). The resolutions of maps of the receptor-bound and unliganded quasi-threefold (q3) spikes were 2.9 Å (masked spike protein trimer) and 3.4 Å (q3 block), respectively (Table S2). In the receptor bound structure, PCDH10 EC1 is inserted into clefts formed by adjacent E2–E1 protomers (Figure 1E). The angle of receptor binding is roughly 45 degrees in the long axis of EC1 with respect to the threefold axis of the spike protein trimer (Figure 1D). PCDH10 EC1 makes extensive contacts with E2 and E1 protomers on both sides of each cleft and buries a large surface area (∼1,500 Å^2^) as it interacts with the WEEV spike protein (Figure S5A). When structures of receptor-bound alphavirus spike proteins are compared, this receptor contact surface is larger than those of alphavirus spike proteins with individual LDLR class A (LA) repeats but smaller than those of human or avian MXRA8 (Figure S5A).^15–22^ There is no conformational change in the WEEV spike protein or in PCDH10 EC1 when structures of the WEEV spike protein bound to EC1 or of EC1 as part of the EC1–EC4 homodimer are compared (Figure S3G).^14^

One face of the cadherin repeat makes prominent contacts with the β-ribbon of E2 domain A (Figure S5D). These contacts involve PCDH10 residues N40 and R42 and WEEV residues D40, D156, and H157, which are in the central arch of the β-ribbon (Figure 1F). Additionally, PCDH10 residues P39, F80, L85, and L87 contact multiple E2 β-ribbon residues including L149, T262, T264, and V265 (Figure 1G). The contralateral face of the cadherin repeat makes several contacts with the adjacent E2–E1 protomer (E2’-E1’) (Figure S5E). As part of these interactions, PCDH10 residues E21 and D22 make salt bridges with E2’ domain B residues K177 and K224 (Figure 1H).

Additionally, PCDH10 residue H76 and E2 residue H21 are involved in stacking interactions (Figure 1I). PCDH10 residues L2 and H3 contact E1’ fusion loop residues F87 and W89 and PCDH10 residue F88 contacts E1’ residue K227 (Figure 1J). Other than the two E1’ fusion loop contact residues, the WEEV spike protein residues that contact PCDH10 are not conserved in EEEV or VEEV, explaining why these other encephalitic alphaviruses do not bind PCDH10 (Figures S6A and S6B).^10^

Interestingly, the hydrophobic interactions the WEEV spike protein makes with human PCDH10 resemble how PCDH10 EC1 interacts with EC4 during antiparallel homodimerization of the receptor ectodomain as part of its physiological function (Figures 1G and 1K).^14^ Therefore, the receptor surface that contacts the WEEV spike protein is occluded in the PCDH10 homodimer, and only the monomeric form of PCDH10 may bind the WEEV spike protein to facilitate viral entry (Figure S3G).

### Assessment of WEEV–PCDH10 interactions

The WEEV CBA87 spike protein contacts fifteen residues on human PCDH10 EC1 (Figure S6C). These PCDH10 residues are highly conserved among PCDH10 orthologs that can serve as WEEV receptors,^10^ which in addition to human PCDH10 includes the murine, equine, avian, and reptilian PCDH10 orthologs. To evaluate the importance of individual interactions of the WEEV spike protein, we performed infectivity assays on K562 cells, a lymphoblast-derived cell line that does not express PCDH10.^10,31^ We transduced K562 cells with a truncated PCDH10 construct containing only wild-type or mutated versions of PCDH10 EC1 as a minimal receptor fragment (Figure 1A). For infectivity assays, we chose the 71V1658 (71V) WEEV strain, a North American strain that binds PCDH10 but not VLDLR or ApoER2 (Figure 1B).^10^

For most of the WEEV-PCDH10 interface, replacing with alanine individual or multiple PCDH10 EC1 residues that contact the WEEV spike protein had no effect on WEEV 71V RVP entry (Figures 1L and S8). However, individually replacing with alanine two EC1 residues that make polar contacts with the central arch of the E2 β-ribbon connector (N40A or R42A) diminished WEEV 71V RVP infection; simultaneous replacement of both residues with alanine completely abolished entry (Figure 1L). Furthermore, the EC1 P39R substitution, which would introduce steric clashes with nearby residues in the E2 β-ribbon connector, completely abrogated WEEV 71V RVP entry (Figure 1L). Thus, in most cases, the large interaction interface between PCDH10 EC1 and the WEEV spike protein is highly tolerant of substitutions that would alter individual contacts with the receptor.

### Structure of WEEV bound to avian PCDH10

The Imperial 181 (lineage B3) WEEV strain was recovered from a mosquito pool in California in 2005 and is nonpathogenic in animal models.^4,24^ Imperial 181 can bind sparrow (*P. domesticus*) PCDH10 but not human PCDH10 as a receptor (Figure 1B).^10^ We used enzyme-linked immunosorbent assays (ELISAs) with VLPs and human and sparrow PCDH10_EC1_–Fc fusion proteins to compare half maximal effective concentrations (EC_50_) values as a surrogate for affinity measurements. As expected, sparrow PCDH10_EC1_–Fc, but not human PCDH10_EC1_–Fc, bound to Imperial 181 VLPs (Figures 2A and S2F). While sparrow PCDH10 _EC1_–Fc bound McMillan (lineage A) and CBA87 (lineage C) VLPs with comparable apparent affinities (∼ 1 nM), human PCDH10 _EC1_–Fc bound both VLPs with vastly different apparent affinities (3.8 µM and 77 nM, respectively) (Figures 2A and S2F). Interestingly, sparrow PCDH10_EC1_–Fc bound Imperial 181 VLPs much less tightly (2.6 µM) than it bound McMillan (1 nM) or CBA87 VLPs (1 nM) (Figures 2A and S2F).

**Figure 2.**
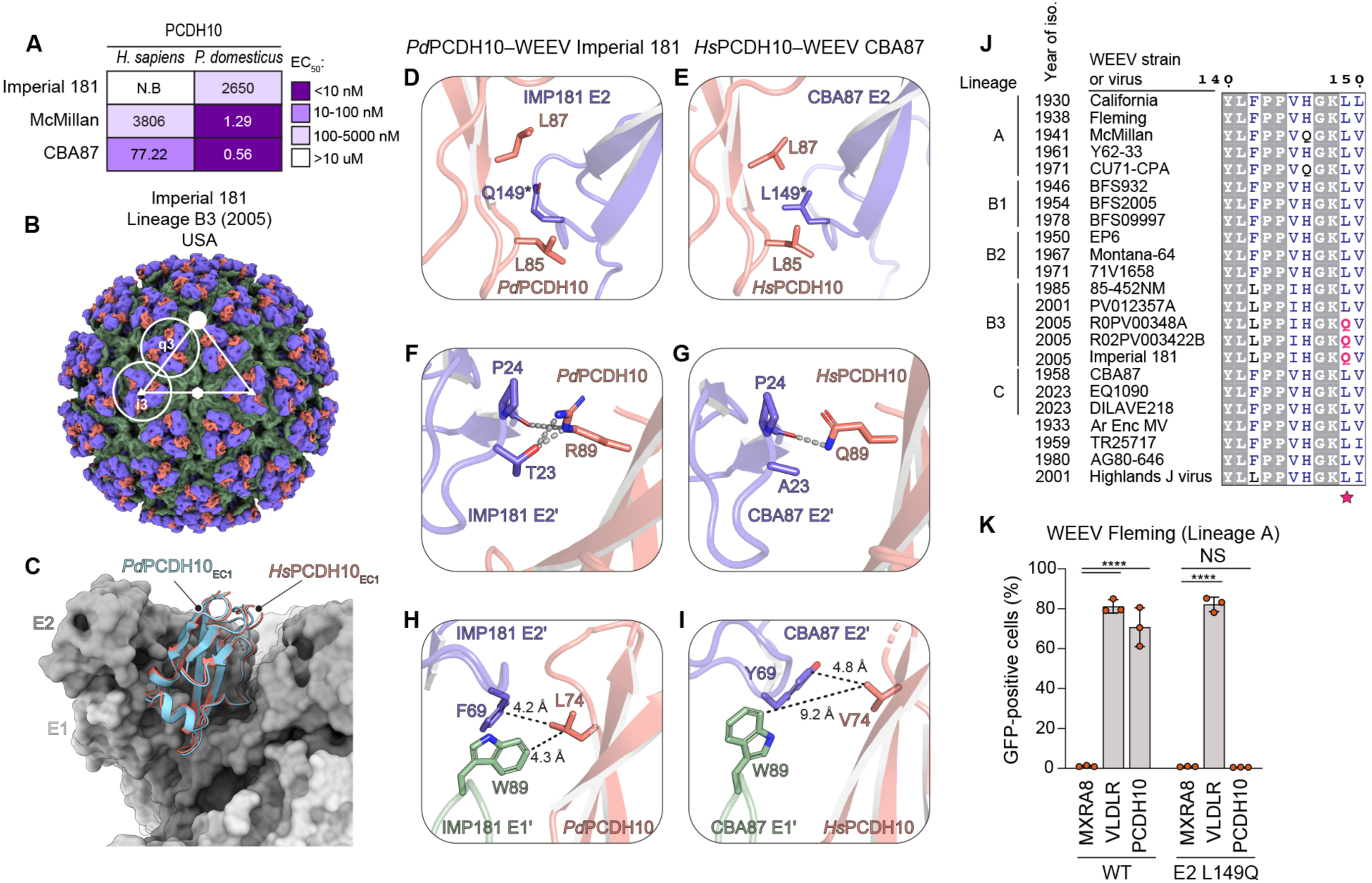
WEEV recognition of avian PCDH10. (A) Summary of the EC_50_ values measured in ELISAs with human (*Hs*) PCDH10_EC1_–Fc or *P. domesticus* (*Pd*) PCDH10_EC1_–Fc on immobilized WEEV VLPs for the indicated strains. See Figure S2F for additional information. (B) Cryo-EM map of *Pd*PCDH10_EC1_–Fc bound to WEEV Imperial 181 VLP. E2, E1, and PCDH10 are colored purple, green, and pink, respectively. (C) Superposition of the cryo-EM structure of *Pd*PCDH10 EC1 (cyan) bound to the WEEV Imperial 181 spike protein and *Hs*PCDH10 EC1 (pink) bound to the CBA87 spike protein. PCDH10 EC1 is shown as a ribbon diagram and the spike protein subunits are shown in dark (E2) and light (E1) gray. (D–I) Side-by-side comparison of the contact residues of *Pd*PCDH10-bound Imperial 181 (D, F, H) and *Hs*PCDH10-bound CBA87 (E, G, I). Residues that participate in interactions are indicated, with polar contacts shown as gray dashed lines. Dashed lines shown in black indicate the closest distances between PCDH10 residue L74 and atoms on the WEEV spike protein for the protomers shown (H, I). The residue at E2 position 149, a key polymorphic residue in the PCDH10 interface, is highlighted with an asterisk. IMP181: Imperial 181. (J) Partial sequence alignment of WEEV E2 proteins, generated using ESPript3.^54^ The key polymorphic residue at position 149 is indicated with a star. Light gray background indicates residues that are completely conserved. Boxed residues indicate positions where a single majority residue or multiple chemically similar residues are present. These residues are shown in dark blue. Glutamine at E2 position 149 (indicated with a star) is colored pink. See also Figures S9A and S10 for additional information. (K) K562 cells stably expressing human MXRA8, VLDLR or PCDH10 were infected with GFP-expressing wild-type or L149Q mutant Fleming RVPs. Infection was quantified by flow cytometry. Data are mean ± s.d. from two experiments performed in duplicates or triplicates (n=3) (K). Two-way ANOVA with Dunnett’s multiple comparisons test, \*\*\*\**P*<0.0001 (K).

To determine why the WEEV Imperial 181 binds sparrow but not human PCDH10, we determined the cryo-EM structure of Imperial 181 VLPs bound to the EC1 repeat of sparrow PCDH10 (Figures 2B, S4B and S7A–D). The global resolution of the map of the receptor-bound (q3) spike was 2.8 Å (masked spike protein trimer) (Table S2). When the human and sparrow PCDH10-bound WEEV structures are compared, the overall binding mode and contact residues are similar (Figures 2C, S7E, and S9A). In the structure of CBA87 bound to human PCDH10, E2 L149 is in a cluster of hydrophobic residues that interacts with PCDH10 EC1; the analogous residue in the E2 protein of Imperial 181, Q149, does not contact EC1, with its polar side chain positioned towards a mostly hydrophobic interface (Figures 2D and 2E). On the receptor side of the interface, the side chain of sparrow PCDH10 residue R89 makes multiple polar contacts with Imperial 181 E2 residues T23 and P24 (Figure 2F). However, in human PCDH10, R89 is replaced by a glutamine (Q89) and the side chain of this residue only contacts the backbone carbonyl of WEEV E2 residue P24, thus providing weaker contributions to the interaction with the receptor (Figure 2G). Additionally, the side chain of residue L74 in sparrow PCDH10 is near a hydrophobic pocket that involves E1’ fusion loop residue W89 (Figure 2H). L74 in sparrow PCDH10 is replaced by a valine in human PCDH10, with the valine side chain positioned further from the pocket and unable to contact E1’ fusion loop residue W89 (Figure 2I).

### An E2 polymorphism drives PCDH10 ortholog dependencies

The leucine residue at WEEV E2 position 149 is highly conserved among WEEV strains in lineage A, B1, B2, and even some B3 strains, and it is only replaced by a glutamine in Imperial 181 and in two additional recent lineage B3 strains that were the most recently isolated in the USA (2005) (Figures 2J, S9A and S10). We had previously tested one of these additional B3 strains (R02PV003422B) and found that it also does not bind human PCDH10 (Figure 1B).^10^ Given that the L149Q substitution would disrupt key hydrophobic contacts WEEV E2 L149 would otherwise make with PCDH10 EC1, we sought to determine whether the leucine at E2 residue position 149 impairs WEEV’s ability to bind human PCDH10. We introduced the E2 L149Q mutation into Fleming (lineage A) RVPs, which can bind human PCDH10, VLDLR, and ApoER2 as receptors.^10^ We used the mutant Fleming RVPs to conduct infectivity assays in K562 cells overexpressing human PCDH10, with human VLDLR or human MXRA8 included as controls. Fleming RVPs containing the E2 L149Q mutation could not infect cells expressing human PCDH10 but could still recognize human VLDLR (Figure 2K). Thus, the L149Q polymorphism likely explains the inability of some B3 strains to bind human PCDH10.

### Structural basis for VLDLR recognition

Lineage A WEEV strains, which were isolated in the 1930–1940s in North America, are highly pathogenic in animal models.^24^ We previously found that lineage A strains, in addition to binding PCDH10, also bind VLDLR and ApoER2 (Figure 1B).^10^ The VLDLR ligand-binding domain (LBD) contains eight LDLR class A (LA) repeats (Figure S12A), which are cysteine-rich domains that contain a Ca^2+^ ion coordinated by multiple acidic residues next to an aromatic residue.^32^ The aromatic residue and neighboring acidic residues usually interact with basic residues on physiological ligands and viruses.^16,17,19–22,33–36^ Importantly, the critical basic residues on the E2 or E1 proteins of alphaviruses that bind to LA repeats are not conserved among alphaviruses (Figures S6A and S6B).^17,19–22,37^ Thus, where LA repeats bind the WEEV spike protein cannot be predicted using previously determined structures.

We first sought to map LA repeat dependencies of WEEV lineage A strain McMillan, a strain that was isolated from a human individual in Canada in 1941. We used K562 cells stably expressing VLDLR truncation constructs in which the LBD is replaced by single LA repeats (Figures S12A and S12B). WEEV McMillan could infect cells expressing single LA repeat constructs of LA1, LA2, LA3, and LA5 (Figures 3A and S12C). Interestingly, the LA repeat binding preferences of WEEV McMillan are different than those described for other alphaviruses (EEEV, Semliki Forest virus, and Sindbis virus).^19–21,37^

**Figure 3.**
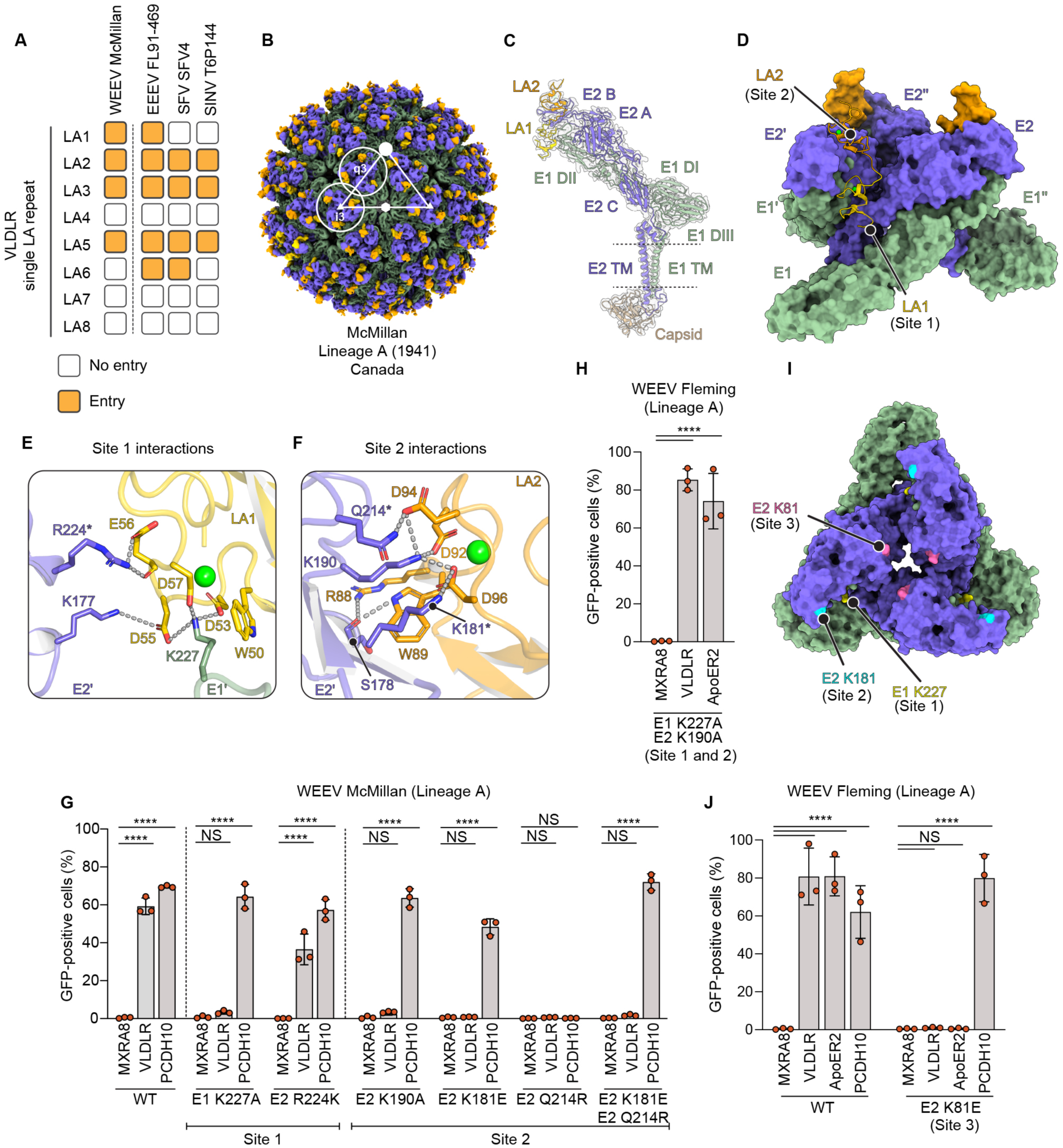
VLDLR recognition by ancestral WEEV strains. (A) Summary of the results of infectivity studies with GFP-expressing RVPs for the indicated alphaviruses with K562 cells expressing VLDLR single LA repeat constructs, with entry quantified using flow cytometry. Infectivity with WEEV McMillan is reported here (see Figures S9A–C) and results for EEEV, SFV, and SINV were previously reported.^19^ “Entry” indicates that the construct mediates statistically significant (p < 0.05) RVP infection when compared to a control construct that lacks a ligand-binding domain (LBD). (B) Cryo-EM map VLDLR_LBD_–Fc bound to WEEV McMillan VLP. E2, E1, and VLDLR LA1 and LA2 are colored in purple, green, light yellow, and dark yellow, respectively. The 5-fold (i5), 3-fold (i3), and 2-fold (i2) icosahedral symmetry axes are indicated respectively with solid circle, triangle, and hexagon. The icosahedral threefold spike (i3) and quasi-threefold spike (q3) are circled and labeled. (C) Ribbon diagram of a single WEEV E2–E1 protomer and LA1–2 from the VLDLR LBD fitted into the cryo-EM density map. E2 and E1 domains are indicated. Dashed lines indicate the position of the viral membrane. TM: transmembrane. (D) Surface representation of the VLDLR-bound WEEV CBA87 spike protein. One of the three VLDLR LA1– 2 segments are shown in ribbon representation. (E and F) Contacts between WEEV McMillan and VLDLR LA1 (E) or LA2 (F). Polar contacts are shown as dashed lines. Ca^2+^ ions are shown as green spheres. Polymorphic residues that contact VLDLR are highlighted with asterisks. (G) K562 cells expressing human MXRA8, VLDLR, or PCDH10 were infected by GFP-expressing wild-type or mutant WEEV McMillan RVPs. (H) K562 cells stably expressing human MXRA8, VLDLR, or ApoER2 were infected with GFP-expressing wild-type or E1 K227A (site 1) + E2 K190A (site 2) mutant WEEV Fleming RVPs. Infection was quantified by flow cytometry. (I) Top view of the WEEV trimeric spike showing three potential binding sites on WEEV E2 and E1 for VLDLR LA repeats. K181 (site 1, cyan) on the E2 glycoprotein and K227 (site 2, yellow) on the E1 glycoprotein are critical contact residues with VLDLR in WEEV McMillan. A lysine at position 81 (site 3, pink) in WEEV Fleming E2 likely binds VLDLR LA repeats. (J) K562 cells stably expressing human MXRA8, VLDLR, or ApoER2 were infected with GFP-expressing wild-type or E2 K81E (site 3) mutant WEEV Fleming RVPs. Infection was quantified by flow cytometry. Data are mean ± s.d. from three experiments performed in triplicates (n=3) (G, H, J). Two-way ANOVA with Dunnett’s multiple comparisons test, \*\*\*\**P*<0.0001 (G, H, I).

We determined the cryo-EM structure of VLDLR_LBD_–Fc-bound to WEEV McMillan q3 spikes at a resolution of 2.9 Å (Figures S4C and S13A–C; Table S2). In the structure of the receptor-bound spike protein, two LA repeats bind clefts and only contact the E2’-E1’ protomer (E2’-E1’) (Figures 3B–D). Our ability to visualize individual side chains in our high-resolution maps allowed us to unambiguously build LA1 and LA2, with LA1 positioned deepest within the cleft (site 1) (Figures 3D and S13D). This binding mode was also consistent with our observation that WEEV RVPs could infect K562 cells expressing VLDLR LA1 or LA2 (Figure 3A).

During interactions with VLDLR LA repeats, LA1 and LA2 bury 746 Å^2^ of surface area on the McMillan spike (Figure S5A). The LA repeat Ca^2+^-coordinating acidic residues usually make critical contacts with one or two basic residues (lysines or arginines) on ligands, while the adjacent LA repeat aromatic residue stacks against the aliphatic portion of the lysine or arginine side chain.^33,34^ In site 1 of the VLDLR-bound WEEV complex, the side chains of LA1 Ca^2+^-coordinating acidic residues encircle WEEV E1 residue K227, and the aliphatic portion of K227 stacks against LA1 W50 (Figure 3E). Acidic residues in LA1 also contact two E2 domain B residues, K177 and R224 (Figures 3E and S5F). On the second LA repeat binding site that is more distal from the viral membrane (site 2), the side chains of LA2 Ca^2+^-coordinating acidic residues interact with K190 on E2’ domain B, with the aliphatic portion of the E2 K190 side chain stacking against W89 in the LA repeat (Figure 3F). Additional polar contacts that involve the side chain or main chain atoms of WEEV E2 residues K181, Q214 and S178, and LA1 residue D94 and R88, further anchor the LA repeat into place. The WEEV McMillan VLDLR binding mode is distinct from how EEEV, SFV, and VEEV interact with LA repeats,^16,17,20,22^ indicating that McMillan independently evolved its solution for binding LA repeats (Figure S14).

We tested the importance of WEEV McMillan-VLDLR contact residues using infectivity assays with wild-type or mutant WEEV McMillan RVPs on K562 cells expressing receptors. E1 residue K227 (site 1) and E2 residue K190 (site 2) are universally conserved in WEEV strains (Figures S9B, S10 and S11). WEEV McMillan RVPs containing either the E1 K227A (site 1) or E2 K190A (site 2) mutations were unable to infect K562 cells expressing VLDLR, despite retaining the ability to infect cells expressing human PCDH10 (Figure 3G). This observation suggests that simultaneous LA repeat engagement of sites 1 and 2 on the WEEV McMillan spike protein is required for efficient VLDLR binding.

All lineage B strains we had previously tested do not bind to VLDLR or ApoER2.^10^ We next tested whether polymorphisms on the spike protein in site 1 (R224K) or site 2 (K181E or Q214R) explain why these strains do not bind VLDLR or ApoER2 (Figure S9B). The E2 R224K substitution when introduced into McMillan RVPs did not prevent infectivity of K562 cells expressing VLDLR, suggesting that lysine and arginine are interchangeable at this position (Figure 3G). However, the K181E or Q214R substitutions when individually introduced into WEEV McMillan RVPs abolished infectivity of K562 cells expressing VLDLR (Figure 3G). These data suggest that K181 and Q214 are required for VLDLR binding by McMillan. The E2 Q214R substitution, in addition to impairing VLDLR binding, unexpectedly also prevented WEEV McMillan RVP infection of cells expressing PCDH10 (Figure 3G), despite E2 residue 214 not being near the PCDH10 binding site. Because R214 can make a salt bridge with E181, as observed in the CBA87 spike protein structure (Figure S13E), we hypothesized that the E2 R214 substitution destabilizes E2 domain B of the WEEV McMillan spike protein by positioning three basic residues near each other (R214, K181, and K190). Consistent with this notion, WEEV McMillan RVPs containing both the K181E and Q214R E2 substitutions, which would allow R214 to make the salt bridge with E181, had the ability to infect K562 cells expressing PCDH10 (Figure 3G).

### Fleming uses a distinct VLDLR binding site

The lineage A strain Fleming has a glutamate at E2 position 181 (site 2), which, based on our mutational analysis, impairs McMillan binding to VLDLR (Figures 3G and S9B). Yet, Fleming can still bind VLDLR (Figure 3J).^10^ In infectivity studies with Fleming mutant RVPs and K562 cells expressing VLDLR, mutation of either E1 residue K227 (site 1) or E2 residue K190 (site 2) had no impact on VLDLR-dependent entry (Figure 3H). We thus examined the sequence of the Fleming spike protein to identify potential binding sites involving unique, surface exposed lysine residues. We identified one such residue near the threefold axis of the Fleming trimeric spike (E2 K81; site 3) that is replaced by glutamate or aspartate in all other strains (Figures 3I and S9B).

Interestingly, E2 K81 is adjacent to another basic residue, K82, that could also be recruited to participate in LA repeat binding in Fleming E2 (Figure S10). Introducing the E2 K81E substitution into Fleming RVPs abrogated WEEV Fleming RVP infection of K562 cells expressing VLDLR or ApoER2, while preserving PCDH10-dependent entry (Figure 3J). Therefore, we propose that the key determinant of LA repeat binding for WEEV Fleming is near the threefold axis of the trimer in a third potential LA repeat binding site in WEEV spike proteins.

### E2 substitutions reinstate WEEV neurotropism

PCDH10 and LDLR-related proteins are expressed on brain cells. Imperial 181 does not bind VLDLR, ApoER2, or PCDH10, and has been shown to replicate poorly in the brain of infected mice,^4,24,38^ suggesting an impaired ability to infect neurons and cause encephalitis as compared to virulent WEEV strains.

To test the hypothesis that Imperial 181’s lack of PCDH10, VLDLR, or ApoER2 binding explains its inability to infect neurons, we generated five mutant Imperial 181 RVPs containing the spike protein substitutions at polymorphic sites that should restore binding to PCDH10, VLDLR and ApoER2, or all three receptors. Mutant 1 (Mut-1) RVPs, which contain the E2 Q149L substitution that should restore the human PCDH10 binding site, were able to infect K562 cells expressing human PCDH10 (Figures 4A–C). Mut-2 RVPs, which contain the E2 E81K substitution that should provide a site 3 LA repeat binding site, could infect K562 cells expressing VLDLR or ApoER2, but not PCDH10. Mut-3 RVPs, which contain the Q149L and the E81K (site 3) substitutions, could infect K562 cells expressing PCDH10, VLDLR, or ApoER2. Mut-4 RVPs, which contain the E181K+R214Q substitutions that would restore a site 2 binding site for LA repeats could infect K562 cells expressing VLDLR and ApoER2. Mut-5 RVPs, containing the Q149L and R214Q+E181K substitutions that would restore PCDH10 binding and a site 2 LA repeat binding site, could infect K562 cells expressing all three receptors.

**Figure 4.**
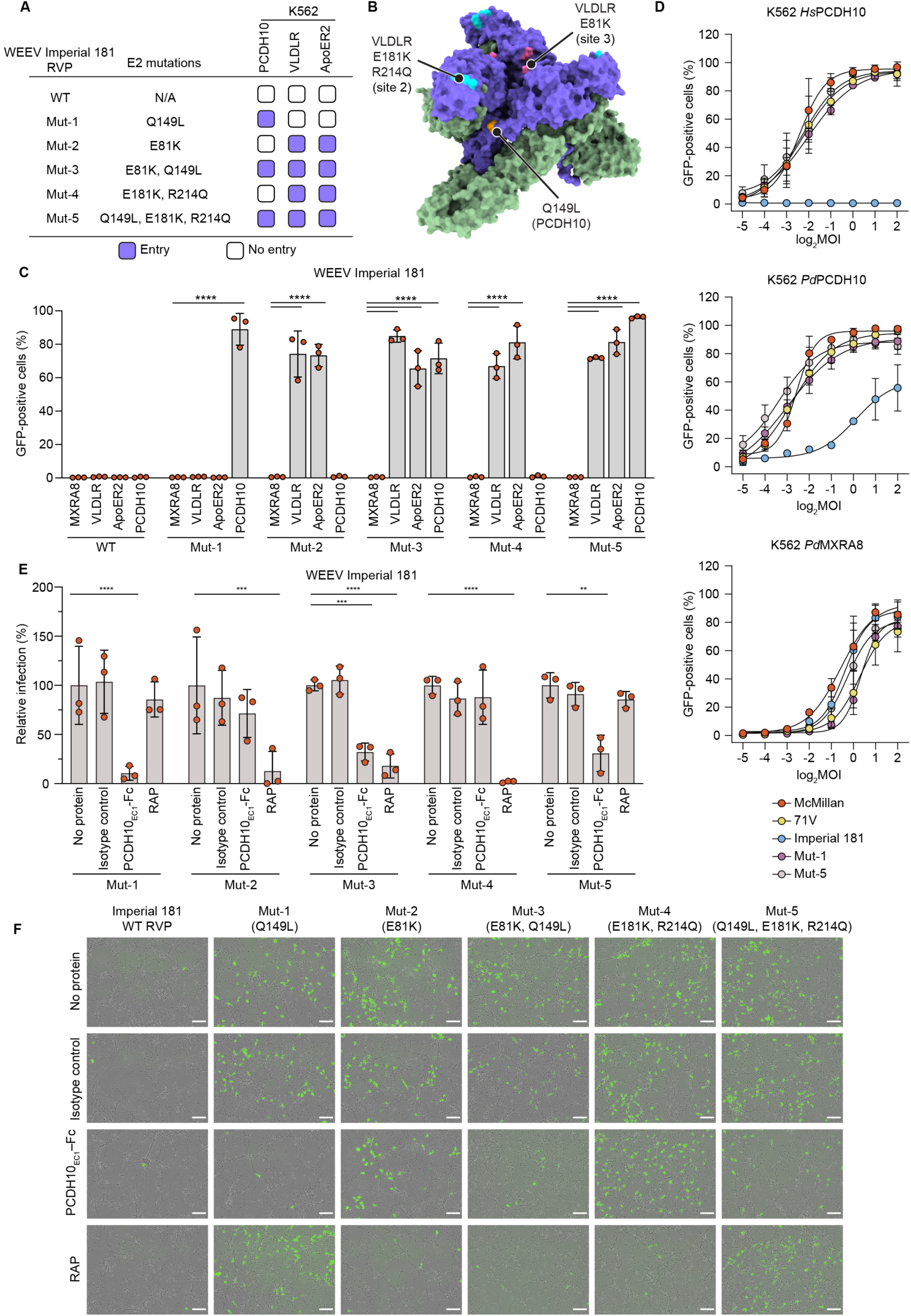
WEEV E2 protein polymorphisms that alter receptor recognition determine WEEV neurotropism. (A) List of mutations generated for WEEV Imperial 181 and summary of K562 infectivity assay as performed in (C). (B) Side view of the WEEV spike highlighting mutated residues on the E2 glycoprotein. (C) K562 cells expressing MXRA8, VLDLR, ApoER2, or PCDH10 were infected with wild-type or mutant Imperial 181 RVPs. Infection was monitored by flow cytometry. (D) K562 cells expressing human PCDH10 (*Hs*PCDH10), sparrow PCDH10 (*Pd*MXRA8), or sparrow MXRA8 (*Pd*MXRA8) were infected with the indicated GFP-expressing RVPs at various MOIs, as pre-determined of Vero E6 cells (see Methods for additional information). Infection was quantified by flow cytometry. (E) Primary murine cortical neurons were infected with GFP-expressing wild-type or mutant WEEV Imperial 181 RVPs in the presence of 316 µg ml^-1^ PCDH10_EC1_–Fc, 100 µg ml^-1^ RAP, or an isotype control. Infection was monitored through a live cell imaging system. (F) Representative merged images of GFP and bright field are shown for the experiment in (E). Scale bars are 100 μm. Data are mean ± s.d. from three experiments performed in duplicates or triplicates (n=3) (C, D, E). One-way ANOVA with Dunnett’s multiple comparisons test, \*\*\*\**P*<0.0001 (C, D). Two-way ANOVA with Šídák’s multiple comparisons test, \*\*\*\**P*<0.0001 (E).

Given the importance of E2 residue L149 in allowing WEEV recognition of human PCDH10, we next assessed the relative efficiency of PCDH10 recognition by mutant Imperial 181 RVPs containing the Q149L substitution in E2. We infected K562 cells expressing human or sparrow PCDH10 with Imperial 181 RVPs at different multiplicities of infection (MOI), with RVP titers determined on Vero E6 cells. We chose Vero E6 cells to measure RVP titers because we previously found that WEEV Imperial 181 RVPs can infect these cells without binding to PCDH10 or LDLR-related proteins.^10^

Imperial 181 RVPs were unable to recognize human PCDH10 to infect K562 cells even at an MOI of 4, but recognized sparrow PCDH10 (Figure 4D). However, Imperial 181 RVPs required a much higher MOI than did McMillan and 71V RVPs to reach 50% infection of cells expressing sparrow PCDH10, suggesting that Imperial 181 uses sparrow PCDH10 less efficiently as a receptor. The results of these infectivity assays are consistent with a lower apparent affinity of Imperial 181 VLPs for sparrow PCDH10_EC1_–Fc as compared to McMillan VLPs in ELISA experiments (Figure 2A). Interestingly, we found that Imperial 181 mutant RVPs that contain the Q149L substitution, Mut-1 and Mut-5, not only gained the ability to infect cells expressing human PCDH10, but also were able to more efficiently infect cells expressing sparrow PCDH10 (Figure 4D). These observations suggest that the L149Q polymorphism decreases affinity for human and sparrow PCDH10, which is likely explained by the removal of key hydrophobic contacts E2 residue L149 makes with PCDH10 EC1 (Figures 2D and 2E). Interestingly, all tested RVPs could infect K562 cells expressing sparrow MXRA8 with similar efficiencies (Figure 4D). This suggests that the E2 L149Q polymorphism does not affect binding to sparrow MXRA8.

We next used the five mutant RVPs to infect primary cortical neurons isolated from embryonic mice. Wild-type Imperial 181 RVPs did not infect murine primary neurons (Figures 4E, 4F, and S12D). However, RVP mutants 1 through 5, consistent with their ability to bind either PCDH10 or VLDLR/ApoER2, all gained the ability to infect murine primary neurons (Figures 4E, 4F, and S12D). Confirming that the infection we observed was specific to the mutant RVPs’ newly acquired ability to bind specific receptors, treatment with PCDH10_EC1_–Fc blocked infection by all mutants that should have gained the ability to bind mammalian PCDH10 because they contain the Q149L substitution (Mut-1, Mut-3, and Mut-5) (Figures 4E and 4F). Treatment with the near-universal LDLR family member ligand antagonist receptor-associated protein (RAP), which blocks alphavirus spike protein binding to VLDLR and ApoER2,^39^ blocked the entry of Imperial 181 mutants that should only bind VLDLR or ApoER2 (Mut-2 and Mut-4) (Figures 4E and 4F). The “Fleming-like” composite Mut-3 was likewise blocked by RAP (Figures 4E and 4F), suggesting that despite containing leucine at E2 position 149 which allows for PCDH10 binding, Mut-3 still critically depends on LDLR-family receptors to infect these neurons. However, the “McMillan-like” composite Mut-5 was not affected by the addition of RAP (Figures 4E and 4F), suggesting that Mut-5, unlike Mut-3, primarily depends on PCDH10 to infect these neurons.

### Spike protein sequence-based prediction of receptor binding patterns

We hypothesized that knowledge of the E2 polymorphisms that drive the ability of WEEV strains to bind human PCDH10 or VLDLR/ApoER2 would allow us to predict the receptor-binding properties of strains we had not yet evaluated. The sequences of nine WEEV strains isolated in the 2023–2024 outbreak in South America have been made publicly available (Table S1).^7^ Phylogenetic analysis places all these strains into a newly designated C lineage (as reported by Campos et al.^7^) that also includes strain CBA87 (Figures 1B and S1). The CBA87 strain was previously isolated from the brain of an infected horse in Argentina in 1958 and is the source strain for the VLPs we used for structural analysis with human PCDH10 EC1.

We included in our analysis three South American WEEV strains that have no lineage assignment (AG80-646, isolated in 1980, Ar Enc MV, isolated in 1933, and TR25717, isolated in Guayana in 1959) (Figure 1B). We also included two re-emerging South American strains, EQ1090 (identified in Brazil in 2023) and DILAVE218 (identified in Uruguay in 2023) (lineage C). All these strains contain a leucine at E2 position 149, which we predict would allow them to bind human and sparrow PCDH10 (Figure 2J). We did not expect these strains to bind VLDLR or ApoER2, as they all lack the lysine residues at E2 positions 181 (site 2) or 81 (site 3) required for VLDLR/ApoER2 binding (Figure S9B). Infectivity assays on K562 cells indeed revealed that RVPs for AG80-646, Ar Enc MV, TR25717, and the two re-emerged lineage C strains could bind PCDH10 but not VLDLR or ApoER2 for cellular entry (Figure 5A). In a second experiment, we transduced K562 cells to express orthologs for nonhuman mammal, avian, and reptilian orthologs that WEEV is known to bind in addition to human orthologs of PCDH10 (Figure 5B). AG80-646 and two lineage C strains we evaluated, CBA87 (1958) and EQ1090 (2023), bound all tested orthologs (Figure 5B).

**Figure 5.**
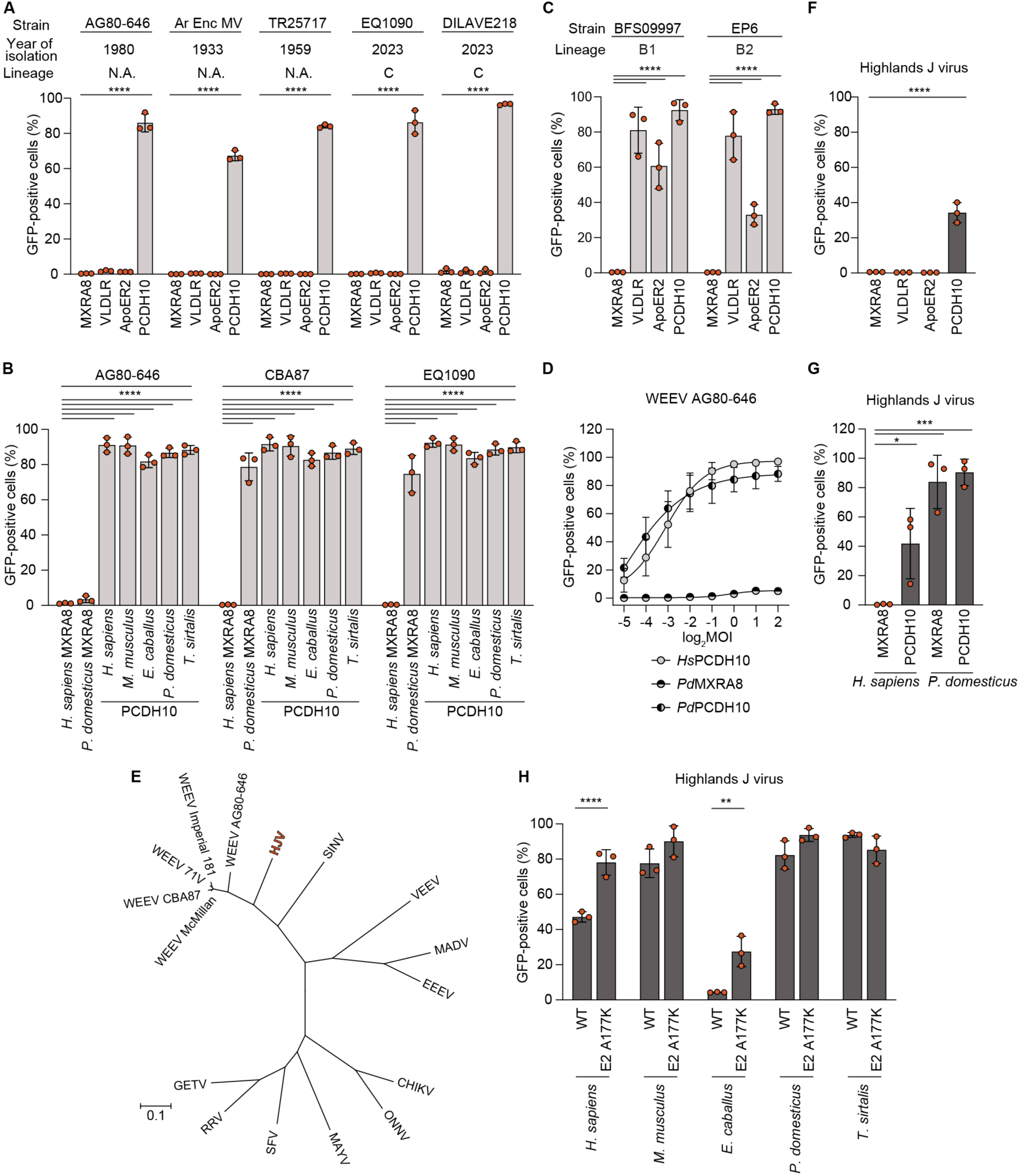
Prediction of alphavirus receptor usage based on spike protein sequences. (A) K562 cells expressing human MXRA8, VLDLR, ApoER2, or PCDH10 were infected with GFP-expressing RVPs for the indicated South American WEEV strains. (B) K562 cells expressing the indicated orthologs of PCDH10 or MXRA8 were infected with GFP-expressing RVPs for the indicated South American WEEV strains. (C) K562 cells expressing human MXRA8, VLDLR, ApoER2, or PCDH10 were infected with GFP-expressing WEEV BFS09997 (lineage B1) and WEEV EP6 (lineage B2) RVPs. (D) K562 cells expressing human PCDH10 (*Hs*PCDH10), sparrow PCDH10 (*Pd*MXRA8), or sparrow MXRA8 (*Pd*MXRA8) were infected with GFP-expressing WEEV AG80-646 RVPs at the indicated MOIs, as determined on Vero E6 cells (see Methods for additional information). (E) Maximum likelihood phylogenetic tree of select alphaviruses (WEEV, highlands J virus (HJV), Sindbis virus (SINV), VEEV, Madariaga virus (MADV), EEEV, Chikungunya virus (CHIKV), O’nyong’nyong virus (ONNV), Mayaro virus (MAYV), Semliki Forest virus (SFV), Ross River virus (RRV), Getah virus (GETV)) using the coding sequences of the structural polyprotein genes. See Table S1 for strain information. (F) K562 cells expressing human MXRA8, VLDLR, ApoER2, or PCDH10 were infected with GFP-expressing HJV RVPs. (G) K562 cells expressing *Hs*PCDH10, *Pd*PCDH10, *Hs*MXRA8, or *Pd*MXRA8 were infected with GFP-expressing HJV RVPs. (H) K562 cells expressing PCDH10 orthologs were infected with GFP-expressing wild-type (WT) and HJV E2 A177K mutant RVPs. This experiment was performed with an MOI of 1 for WT and mutant RVPs measured on Vero E6 cells. Infection was quantified by flow cytometry. Data are mean ± s.d. from three experiments performed in duplicates or triplicates (n=3) (A, B, C, D, F, G, and H). Two-way ANOVA with Dunnett’s multiple comparisons test (A, B, C). Two-way ANOVA with Šídák’s multiple comparisons test (H). One-way ANOVA with Dunnett’s multiple comparisons test (F, G)

When we examined polymorphisms in VLDLR binding sites across North American WEEV strains, we found that two lineage B strains we had not previously tested, BFS09997 (sublineage B1, isolated in 1978 from mosquitoes in California, USA) and EP6 (sublineage B2, isolated in 1950 from mosquitoes in Missouri, USA), contain a lysine at E2 position 181 (LA repeat binding site 2) (Figure S9B), which may allow these strains to bind VLDLR and ApoER2. Unlike all other lineage B strains we had previously tested, BFS09997 and EP6 RVPs could infect K562 cells that express VLDLR and ApoER2 (Figure 5C). Interestingly, both strains have R214 (Figure S9B), which our mutagenesis studies suggested prevents VLDLR/ApoER2 binding in the context of the McMillan spike protein (Figure 3G). Thus, these two lineage B strains can tolerate the combination of E2 K181 and R214, which could be related to compensatory changes elsewhere in their spike proteins. Thus, our data suggest that E2 K181 is a determinant of VLDLR and ApoER2 binding that can be acquired by strains outside of the ancestral A lineage.

### Shift in receptors of a South American WEEV strain

AG80-646 is an enzootic WEEV strain that was isolated from mosquitoes in Argentina in 1980,^40^ and likely represents a lineage that is highly divergent from lineage C (Figures 1B and S1). While all WEEV strains we and others have examined to date can bind avian orthologs of MXRA8,^10,41^ unexpectedly, we found that AG80-646 RVPs could not recognize sparrow MXRA8 to infect K562 cells, despite the ability to recognize human, murine, equine, sparrow, and reptilian PCDH10 (Figure 5B). Infectivity studies with AG80-646 RVPs at a range of MOIs on K562 cells expressing sparrow MXRA8 suggests that this strain does not have any affinity for sparrow MXRA8 (Figure 5D). This observation suggests that like WEEV lineages in North America, South American WEEV lineages may have also undergone shifts in their receptor-binding properties as they diverged.

### A WEEV-related alphavirus binds PCDH10

Highlands J virus (HJV) is a North American alphavirus closely related to WEEV that was first isolated in 1960 from a blue jay (*Cyanocitta cristata*) in Florida, USA (Figures 5E and S1; Table S1). HJV is thought to circulate on the East Coast of the USA,^2^ with birds as a primary reservoir, and was most recently detected in a red-tail hawk (*Buteo jamaicensis*) in Georgia in 2001^42^ and in a sandhill crane in *Missisipi* (*Grus canadensis pulla*) in 2012.^43^ The virus is pathogenic in certain avian species.^44,45^ Although cases of human HJV infections have been documented incidentally in individuals co-infected with St. Louis encephalitis virus,^46^ and HJV was associated with a case of fatal equine encephalitis,^47^ HJV is not generally considered to be an equine or human pathogen. HJV has no known receptors.

Conservation of PCDH10 contact residues in the HJV E2–E1 spike protein sequence, including leucine at a position equivalent to WEEV E2 residue 149, led us to suspect that PCDH10 may serve as a receptor for HJV (Figure S9A). Indeed, we found that HJV RVPs could infect cells expressing human, murine, avian, and reptilian orthologs of PCDH10 (Figures 5F–H). Consistent with the lack of the required lysine residues in positions equivalent with WEEV E2 LA repeat binding sites 2 (e.g., K181) or 3 (e.g., K81), HJV could not use VLDLR or ApoER2 to infect cells (Figures 5F and S9B). Interestingly, we also found that avian MXRA8 could serve as a receptor for HJV, consistent with a close evolutionary relationship between HJV with WEEV (Figures 5E and 5G).

Unexpectedly, the HJV strain we tested (585-01, isolated in 2001) could not efficiently enter cells expressing equine PCDH10 (Figure 5H). Examination of sequence alignments and the structures revealed that a K177A polymorphism (with respect to WEEV E2 sequence) found in HJV would remove a key salt bridge with PCDH10 (Figures 1H and S9A). The A177K E2 substitution, when introduced into the HJV RVPs, allowed these RVPs to more readily infect K562 cells expressing equine PCDH10 (Figure 5H). Thus, HJV is another North American alphavirus that can bind PCDH10 orthologs as cellular receptors.

## Discussion

We report here structures of WEEV bound to two of its structurally unrelated alternate receptors, PCDH10 and VLDLR. The structures allow us to define the polymorphisms in the WEEV E2 spike protein that drive shifts in the receptor-binding properties of strains isolated over the past century.

We constructed a phylogenetic tree using the structural polyprotein coding sequences of 57 WEEV strains (Figure S1). The phylogenetic tree shows that E2 Q149, which abolishes human PCDH10 binding, is a signature substitution of the most recently isolated North American B3 clade, to which Imperial 181 belongs (Figure S1). Our tree also shows that newer isolates of North American WEEV strains tend to be placed in new clades instead of extending older clades, confirming previous findings of rapid clade displacement in circulating WEEV.^4,48^ If clade displacement has continued in the past two decades, the E2 Q149 clade may have displaced older clades and expanded in the continent, marking the epizootic decline of WEEV in North America as function of the inability of these newer WEEV strains to bind human and equine orthologs of PCDH10. Indeed, strains in this clade were isolated in California and Texas (Table S1), suggestive of its geographic expansion. Further environmental sampling is required to test this hypothesis.

Strains McMillan and California appear to form a distinct clade within lineage A; this clade can be defined as containing a lysine at E2 position 181. We found that K181 is a contact residue for LA repeats and is a critical determinant of VLDLR and ApoER2 binding (Figures 3F and 3G). This lysine, however, is likely replaced by a glutamate in the common ancestor of WEEV lineage A, given that Fleming contains a glutamate at that position and lies basal to California and McMillan in the phylogenetic tree (Figure S1). Therefore, the clade containing California and McMillan likely evolved from an ancestral strain containing a glutamate at E2 position 181 that acquired a lysine substitution; this ancestral strain went on to spill over and gave rise to large epidemics that spanned over at least a decade in the 1930s–1940s. The McMillan/California clade would later become extinct and displaced. Our identification of two lineage B strains that contain the E2 E181K polymorphism and can bind VLDLR and ApoER2 in addition to PCDH10 (BFS09997 and EP6) (Figure 5C) supports the notion that this polymorphism may be a transitory mutation that could arise stochastically. Of note, while the lineage A strains McMillan and California were more extensively passaged, BFS09997 and EP6 were minimally passaged (Table S1), suggesting that E2 K181 is a naturally occurring substitution as opposed to the result of passaging.

Our mutational analysis suggests that the Fleming strain uses a different surface to bind VLDLR and ApoER2. This strain likely represents a separate lineage A clade that circulated at the same time as strains California and McMillan. As an alternative explanation, a previous study proposed that the E2 K81 polymorphism in Fleming may be a result of cell culture adaptation during passaging.^49^ Strain Mn548, a group B2 strain also containing E2 K81 (Figures S1 and S9B), has unknown source of isolation and passage history, making it difficult to assess the origin of the E2 K81 polymorphism. Nonetheless, our finding that the E81K polymorphism allows WEEV strains to bind VLDLR and ApoER2 to infect neurons suggest that this additional potential site of LA repeat binding would be important to monitor during environmental viral sampling efforts.

Strains from the South American C lineage, which is responsible for the recent re-emergence of WEEV in South America,^7^ all contain a leucine at E2 position 149. The leucine at this position should allow isolated strains to bind human and equine PCDH10 in addition to avian PCDH10. While WEEV has submerged as a human pathogen in North America, in contrast, in South America, serological evidence suggests that WEEV continued to spill over into epizootic hosts well into the 21^st^ century.^50–52^ The ability of WEEV to persist in epizootic circulation in South America may explain the preservation of E2 L149, as well as mammalian PCDH10 recognition, by lineage C in South America.

Our structural analysis of PCDH10- or VLDLR-bound WEEV reveals an overlapping binding site for multiple structurally unrelated receptors. It also explains how receptor-decoy proteins comprising the EC1 domain of PCDH10 or the VLDLR LBD can block access to PCDH10, VLDLR, and ApoER2, and protect mice from lethal challenge by the highly virulent lineage A strain McMillan^10,23^. Additionally, receptor decoy proteins that contain VLDLR LA1-LA2, which are known to be protective against EEEV challenge in mice,^20^ would presumably also protect against WEEV strains that bind LDLR-related proteins.

Interestingly, the EC1 residues of PCDH10 that interact with WEEV are not conserved in its two closest δ2 protocadherin family members, PCDH17 and PCDH19, suggesting that WEEV cannot infect cells by binding to PCDH17 or PCDH19 (Figure S6D).^14^ Although we used a construct that contains only EC1 for structural analysis, the PCDH10 ectodomain includes six cadherin repeats and is thought to be relatively rigid, as calcium ions are coordinated by adjacent repeats. Superimposing the X-ray crystal structure of the PCDH10 ectodomain containing EC repeats 1 through 6^14^ with our structure of WEEV bound to PCDH10 EC1 revealed steric clash at the EC2 domain if all three clefts of the trimeric WEEV spike are bound by the receptor (Figure S3H). We thus suspect that each WEEV trimeric spike may engage only one PCDH10 molecule.

The WEEV PCDH10 binding mode may thus parallel how rhinovirus-C interacts with cadherin-related protein 3 (CDHR3).^53^ In the CDHR3–rhinovirus-C interaction, the most membrane distal repeat of CDHR3 (EC1) also interacts with the virus but the subsequent repeat (EC2) limits occupancy through steric hindrance.^53^

A previous study by Mossel et al. examining the effects of WEEV E2 polymorphism on pathogenicity in mice showed that introducing the E2 Q214R mutation, which our assays revealed would prevent McMillan from binding PCDH10, VLDLR, or ApoER2 (Figure 3G), decreases McMillan’s mortality in mice (from 100% with the wild-type virus down to 10% with the Q214R McMillan mutant) (Figure S9C). Introducing the K181E mutation in WEEV McMillan, which our assays revealed would impair binding to VLDLR and ApoER2 but would still allow for PCDH10 binding (Figure 3G), had a milder effect in decreasing mortality in mice to 70%, while delaying the mean time to death by two days (Figure S9C).^38^ Thus, the ability to bind PCDH10, VLDLR, and ApoER2 likely influences WEEV pathogenicity in mice. The structural information provided in our study will facilitate additional studies that directly evaluate the individual contribution of WEEV mammalian receptors to neurovirulence in animal models.

Our findings provide a guide for estimating the potential threat any given WEEV strain might pose based on sequence polymorphisms in its viral spike protein as determinants of virus receptor dependencies and propensities for neurotropism. They should thus facilitate environmental surveillance and bolster future outbreak preparedness.

**Supplementary Figure 1.**
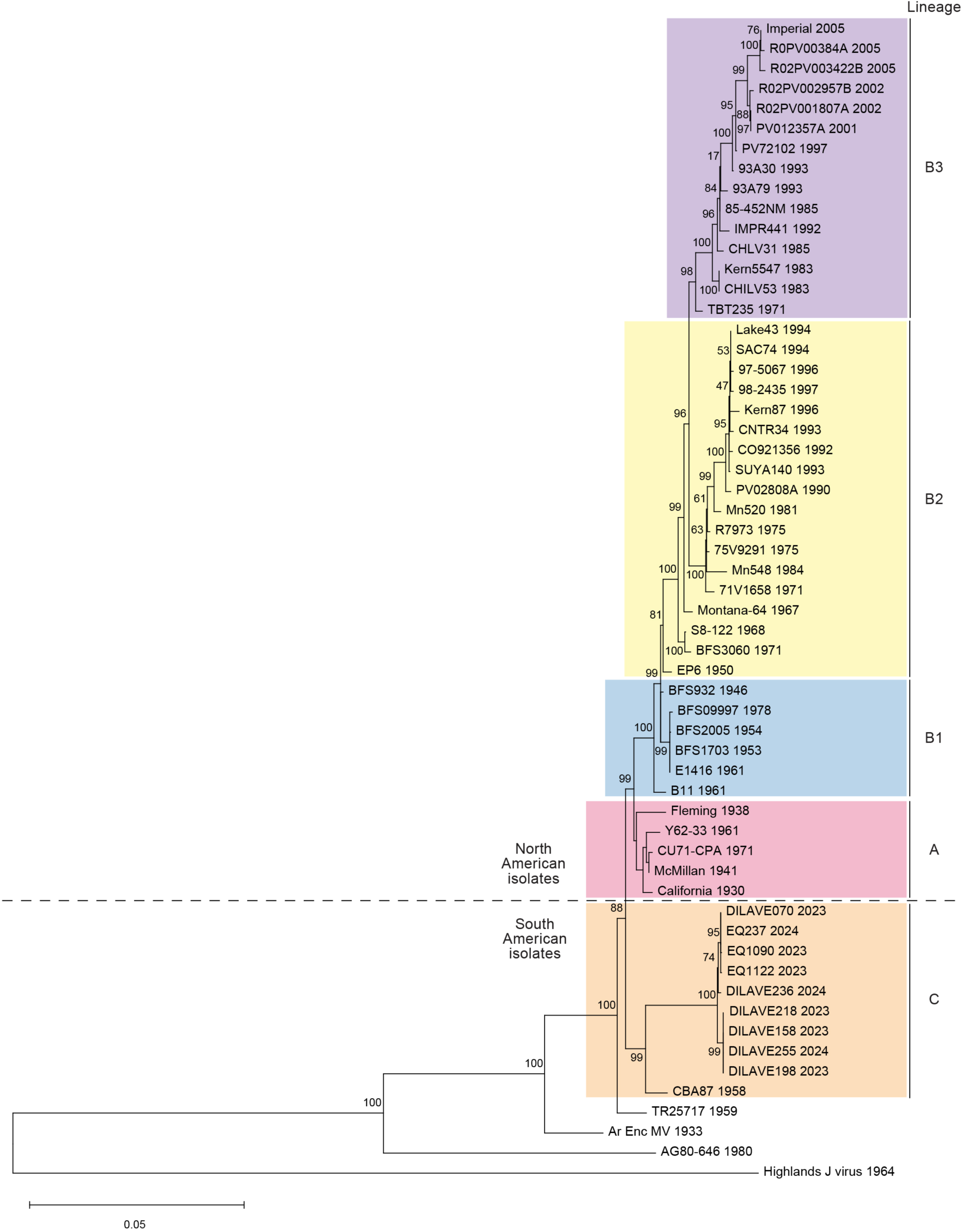
Phylogenetic tree of WEEV strains and Highlands J virus. Maximum likelihood phylogenetic tree of 57 WEEV strains using the coding sequences of the structural polyprotein genes. Highlands J virus strain 64A-1519 (GenBank: KT429021) is also included. Scale bar represents 0.05 nucleotide substitutions per site. SFV strain SFV4, VEEV strain TC-83, and MADV strain 267113 were included in the phylogenetic analysis (not shown). Numbers at nodes indicate bootstrap values. In cases in which the branches are too small, bootstrap values may not be shown. The three lineages (A, B, and C) and B sublineages (B1, B2, and B3) are indicated. Taxon labels include strain name and year of isolation. GenBank accession numbers are provided in Table S1.

**Supplementary Figure 2.**
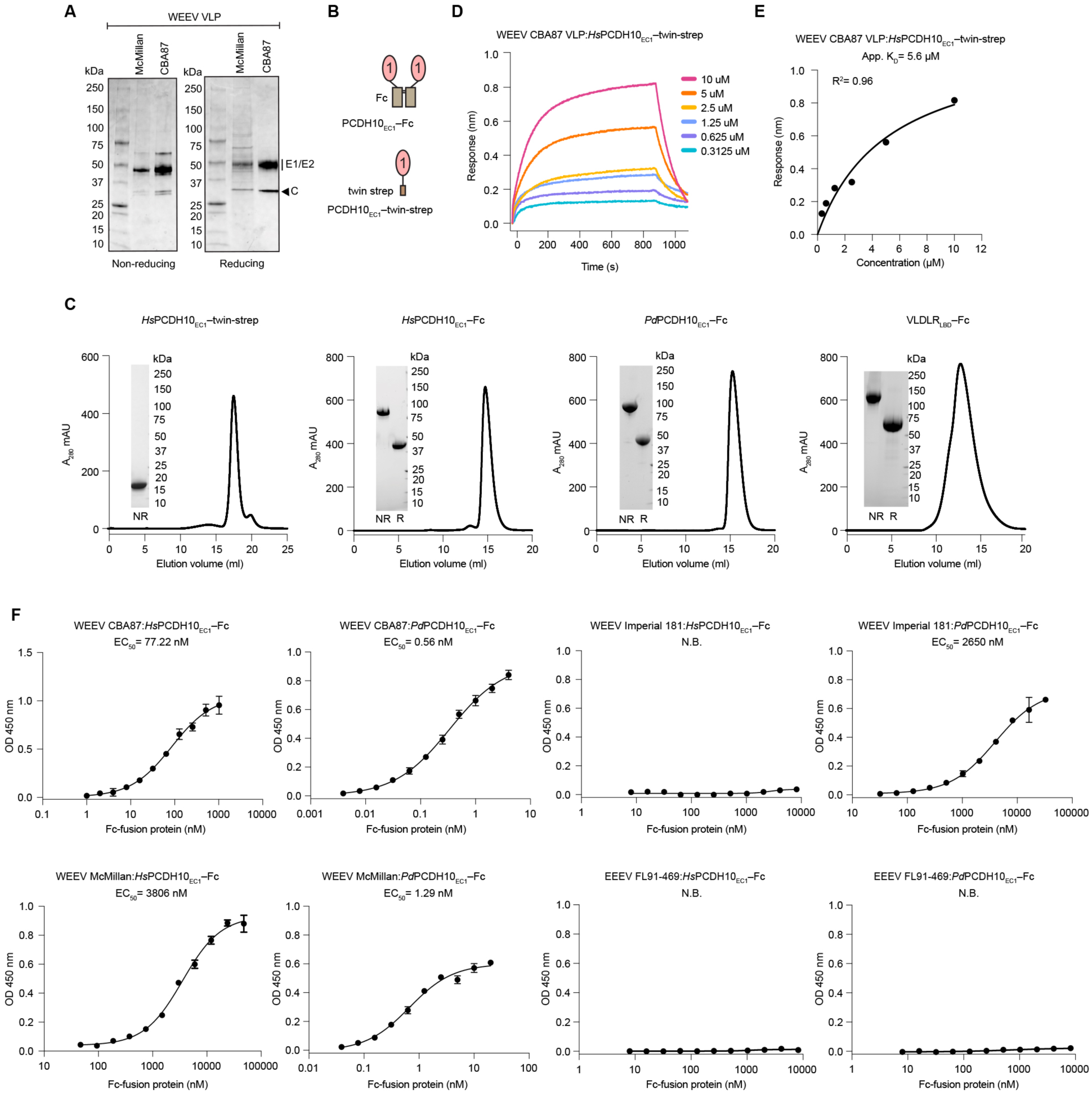
Binding experiments with PCDH10 EC1 and WEEV VLPs. (A) Coomassie-stained SDS-PAGE gel of purified VLPs. The experiment was performed twice, and representative gel images are shown. (B) Schematic diagrams of the PCDH10_EC1_–Fc and the PCDH10_EC1_ twin-strep tag constructs. (C) Size exclusion chromatography of Fc-fusion and twin-strep tagged proteins. Insets are SDS-PAGE gels of pooled peak fractions, which were visualized using a stain-free imaging system.(D) Biolayer interferometry binding analysis of *Hs*PCDH10_EC1_– twin-strep with immobilized WEEV CBA87 VLPs. (E) Scatchard plot for the binding data of *Hs*PCDH10_EC1_–twin-strep with WEEV CBA87 VLPs shown in (D). The measured apparent affinity is indicated. (F) Representative ELISA results showing the binding of the *Hs*PCDH10_EC1_–Fc or *Pd*PCDH10_EC1_–Fc to WEEV CBA87, WEEV McMillan, WEEV Imperial 181, or EEEV FL91-469 VLPs. The experiment was performed three times and representative data are shown. EC_50_ values shown are the mean of three independent experiments.

**Supplementary Figure 3.**
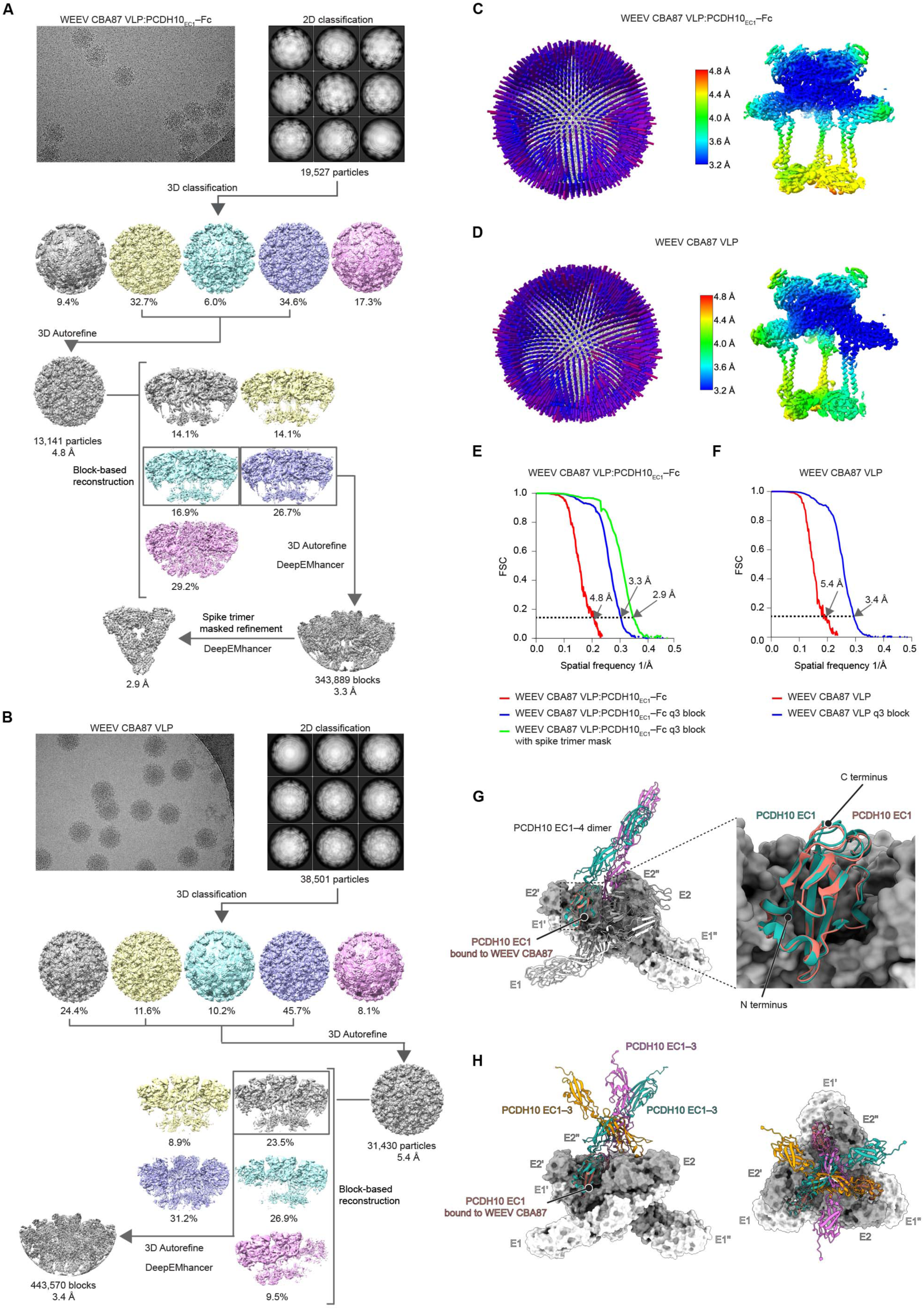
Cryo-EM reconstructions of WEEV CBA87 VLPs alone or bound to human PCDH10_EC1_–Fc. (A) Workflow used for cryo-EM data processing of WEEV CBA87 VLPs bound to human PCDH10_EC1_–Fc. (B) Workflow used for cryo-EM data processing of unliganded WEEV CBA87 VLPs. See Methods for additional details. (C and D) 3D representation of particles angular distribution and local resolution map of the WEEV CBA87 VLP alone (D) or in complex with PCDH10_EC1_–Fc (C). (E and F) Fourier shell correlation curves of WEEV CBA87 VLP alone (F) and bound to PCDH10_EC1_–Fc (E) are shown, respectively. The threshold used to estimate the resolution is 0.143. (G) Structural superimposition of the X-ray crystal structure of PCDH10 EC1–4 homodimer (PDB ID: 6VFW)^14^ with the WEEV CBA87:PCDH10_EC1_–Fc complex (left). (H) Superposition of the X-ray crystal structure of PCDH10 EC1–6 (PDB ID: 6VG4)^14^ onto the EC1 protomers bound to trimeric spike of WEEV CBA87. Occupancy of the three receptor-binding sites on the trimer would result in showing steric hindrance between the EC2 repeat of neighboring receptor molecules. The left panel shows a side view, and the right panel shows a top view.

**Supplementary Figure 4.**
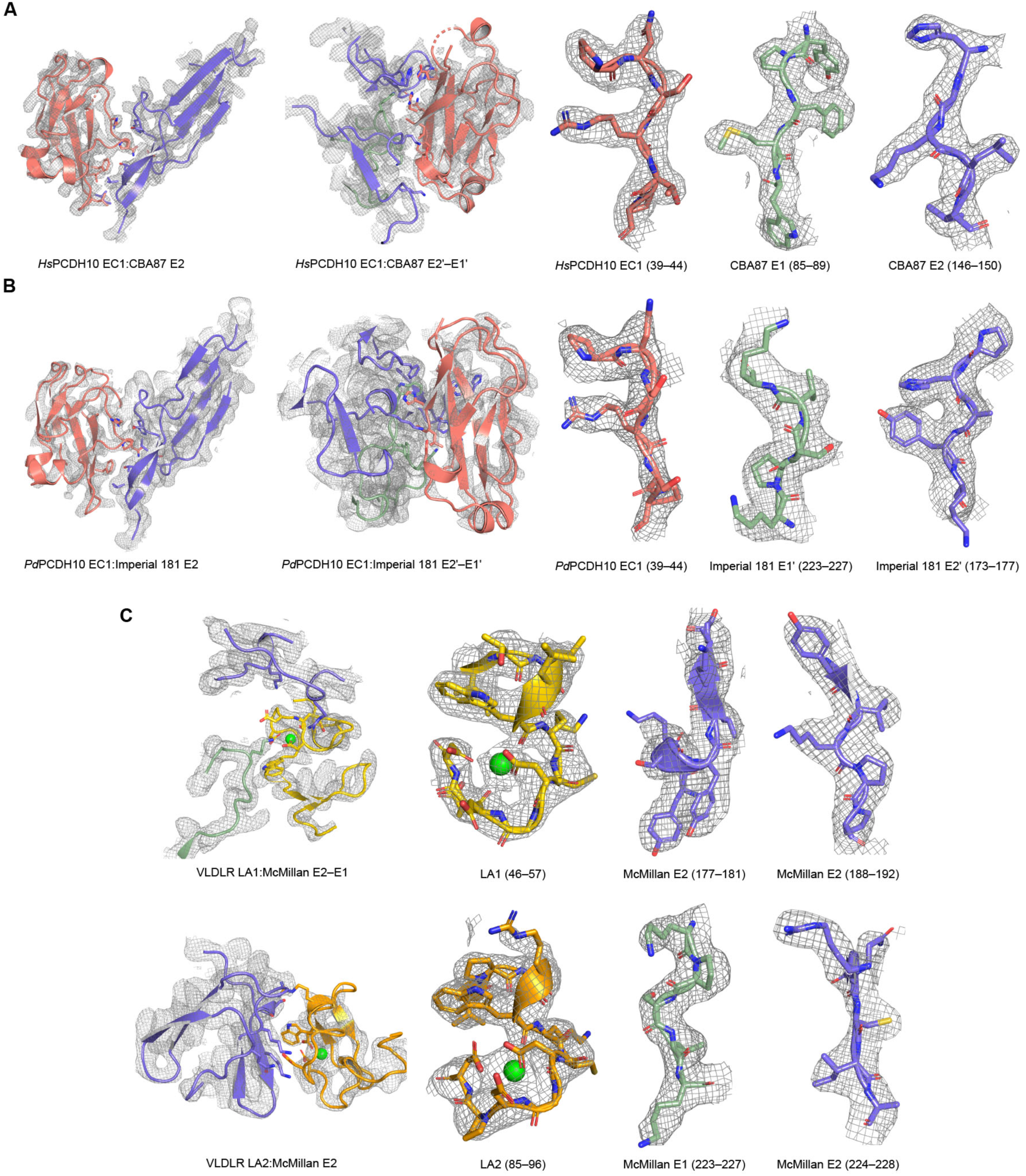
Representative cryo-EM density maps of WEEV VLPs bound to *Hs*PCDH10, *Pd*PCDH10, and VLDLR. (A–C) Density maps of the indicated polypeptide segments from the following complexes: (A) WEEV CBA87 VLP bound to *Hs*PCDH10_EC1_–Fc, (B) WEEV Imperial 181 VLP bound to *Pd*PCDH10_EC1_–Fc, and (C) WEEV McMillan VLP bound to VLDLR_LBD_–Fc.

**Supplementary Figure 5.**
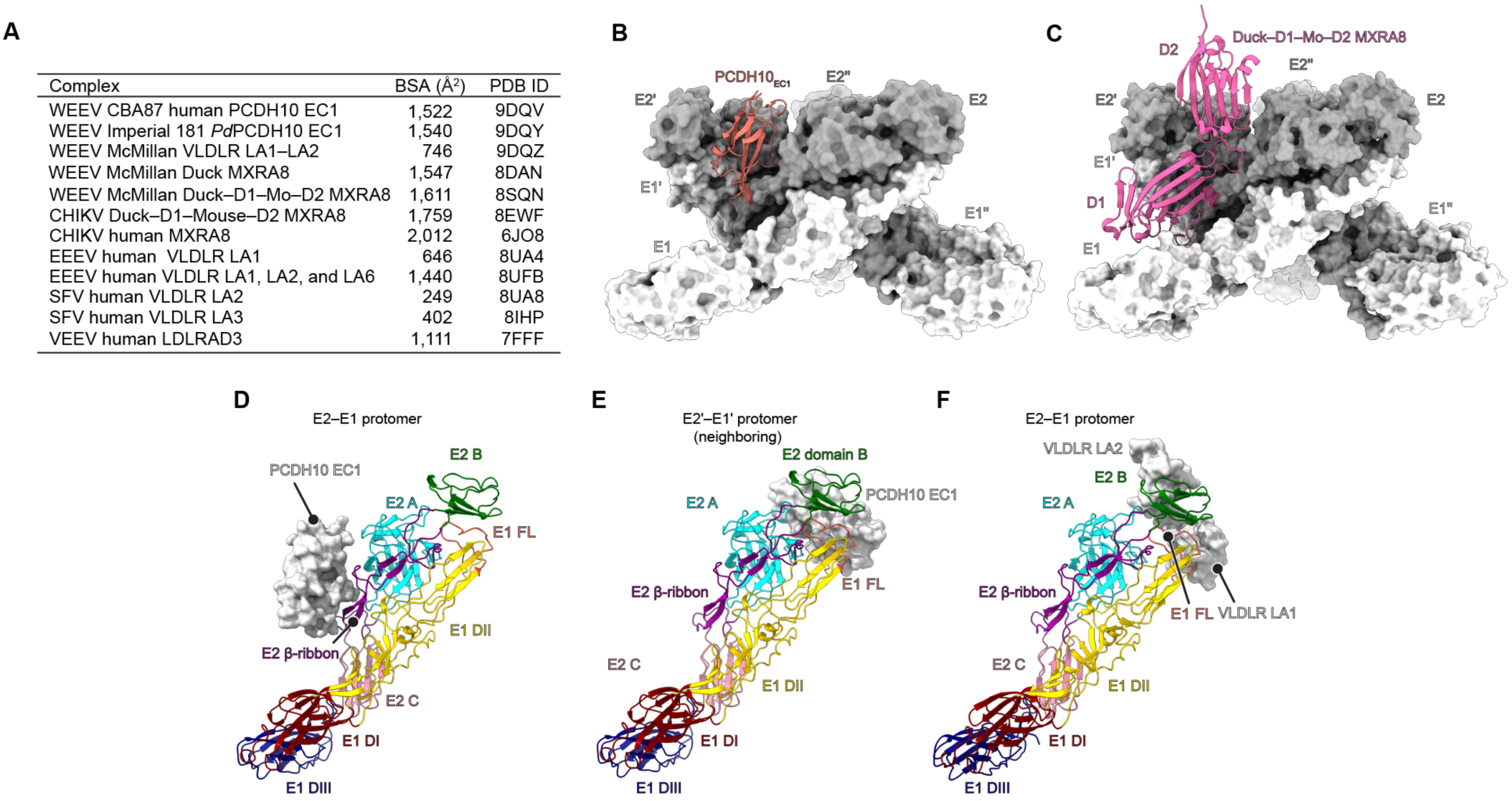
Comparison of contact surfaces for alphavirus E2 and E1 in receptor binding. (A) Buried surface area (BSA) calculations for the indicated complexes. (B) Surface rendering of the WEEV CBA87 spike protein trimer with one protomer of PCDH10 EC1 shown in ribbons. (C) Surface rendering of the WEEV McMillan spike protein with one protomer of chimeric Duck–D1–Mouse–D2 MXRA8 is shown in ribbons (PDB ID: 8SQN).^41^ (D–F) Ribbon diagrams of WEEV E2–E1 heterodimers bound to surface-rendered PCDH10 EC1 or VLDLR LA1–LA2. E2 domains (A, B, and C) and E1 domains (DI-III) are indicated in different colors. Domain organization is shown as originally described by Voss et al.^55^ FL: fusion loop.

**Supplementary Figure 6.**
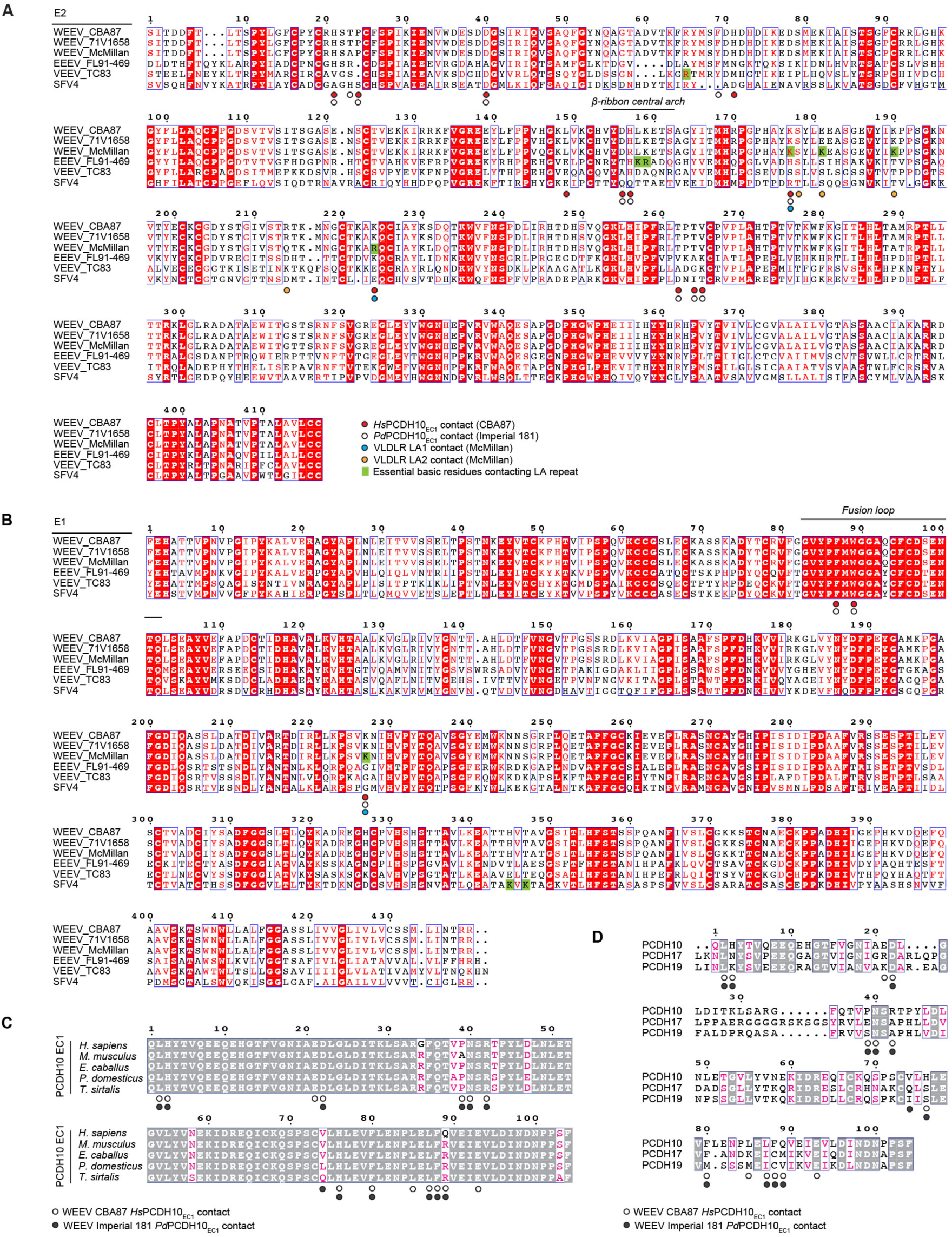
Sequence alignments of alphavirus E2–E1 glycoproteins and protocadherin EC1 repeats. (A and B) Sequence alignments of the E2 (A) and E1 (B) glycoproteins of WEEV CBA87 (GenBank: DQ432026.1), WEEV 71V658 (GenBank: NP_640331.1), WEEV McMillan (GenBank: DQ393792.1), EEEV FL91-469 (GenBank: Q4QXJ7.1), VEEV TC83 (GenBank: AAB02517.1), and SFV4 (GenBank: AKC01668.1). E2 and E1 residues that are with 4 Å of the indicated receptors in the structures are shown as indicated in the legend, and essential basic residues contacting LA repeats based on prior structural studies of LA repeat bound alphavirus spike proteins^17,19–22,37^ are highlighted in green. Residues that are completely conserved in all aligned sequences have a red background. Boxed residues show positions where a single majority residue or multiple chemically similar residues are found. Such residues are in red. (C) Sequence alignments of PCDH10 EC1 orthologs of *H. sapiens* PCDH10 (GenBank: NP_116586.1); *M. musculus* PCDH10 (GenBank: NP_001091642.1); *E. caballus* PCDH10 (GenBank: XP_023492316.1); *P. domesticus* PCDH10 (GenBank: XP_064272564.1); *T. sirtalis* PCDH10 (GenBank: XP_013928164.1). (D) Sequence alignments of human PCDH10 EC1 with the EC1 repeats of other non-clustered δ2 protocadherins PCDH17 (GenBank: NP_001035519.1) and PCDH19 (GenBank: NP_001171809.1). Residues that are completely conserved have a gray background. Boxed residues highlight positions where a single majority residue or multiple chemically similar residues could be identified. Such residues are highlighted in pink. The panels were generated using ESPript 3.0.^54^ PCDH10 residues that are with 4 Å of the indicated spike proteins in the structures are shown as indicated in the legend.

**Supplementary Figure 7.**
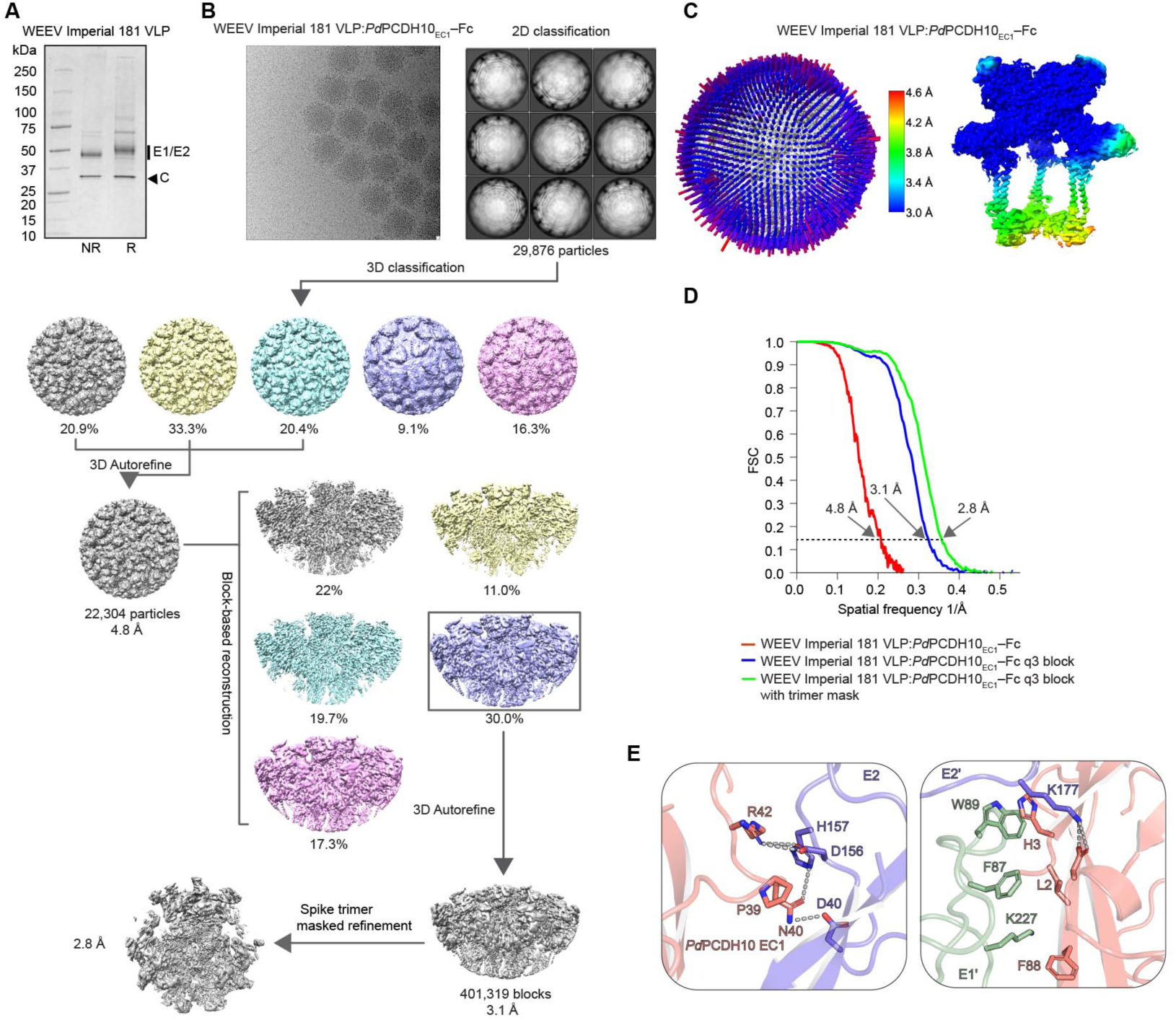
Cryo-EM reconstruction of WEEV Imperial 181 VLP in complex with *Pd*PCDH10_EC1_–Fc. (A) Coomassie-stained SDS-PAGE gel of purified WEEV Imperial 181 VLPs. NR: non-reducing. R: reducing. (B) Workflow used for cryo-EM data processing of WEEV Imperial 181 VLPs bound to *Pd*PCDH10_EC1_–Fc. (C) 3D representation of particles angular distribution and local resolution maps of WEEV Imperial 181 VLP in complex with *Pd*PCDH10_EC1_– Fc. (D) Fourier shell correlation curves of WEEV Imperial 181 VLP in complex with *Pd*PCDH10_EC1_–Fc. The threshold used to estimate the resolution is 0.143. See Methods for additional details. (E) Interface between WEEV Imperial 181 E2–E1 or E2’–E1’ and *Pd*PCDH10 EC1. Residues that participate in interactions between WEEV Imperial 181 E2–E1 heterodimers and *Pd*PCDH10 EC1 are indicated, with polar contacts shown as dashed lines.

**Supplementary Figure 8.**
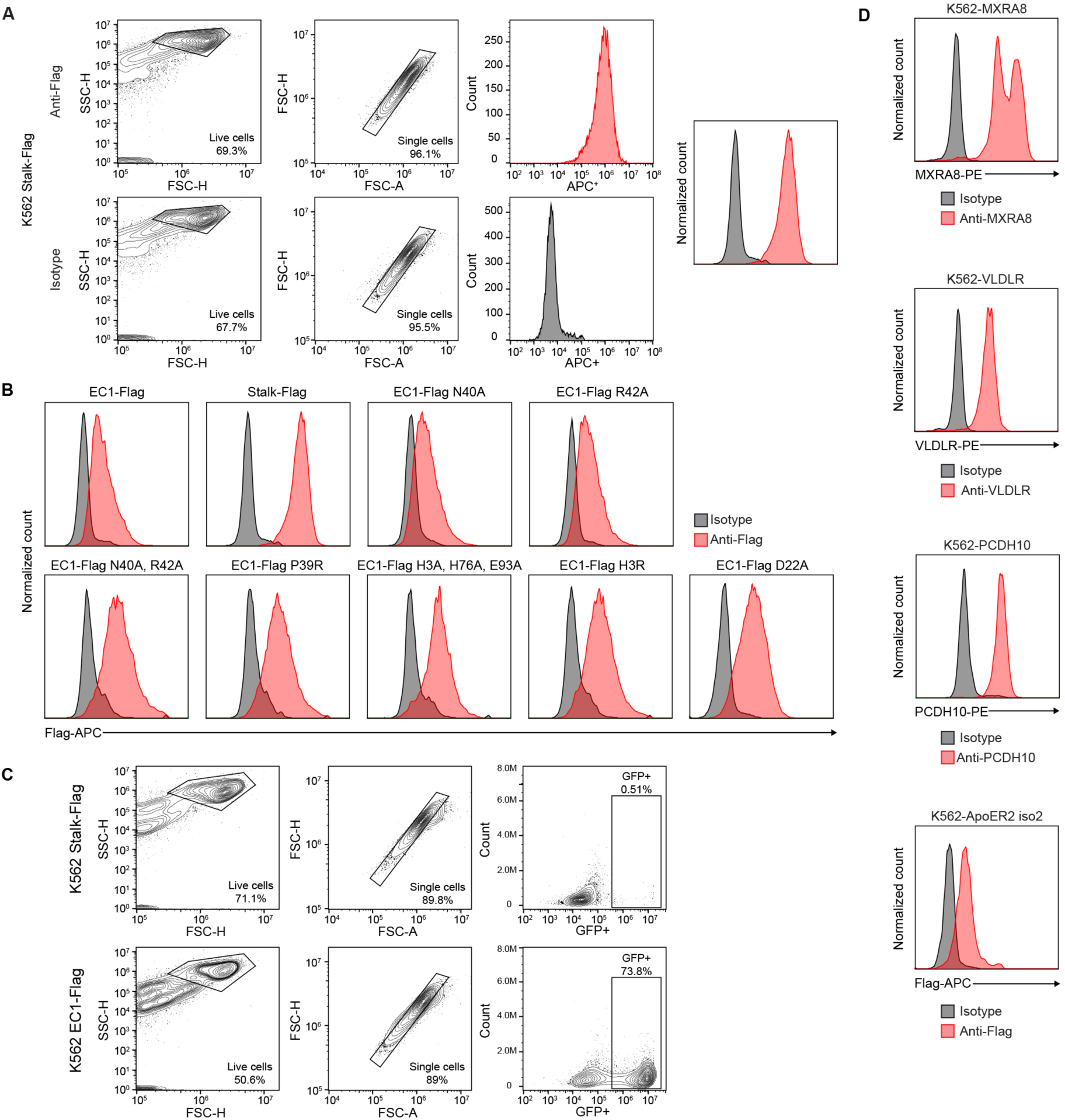
Gating strategy and cell surface receptor staining. (A) Example of flow cytometry gating strategy used to quantify cells stained by antibodies. (B) Staining of K562 cell surface PCDH10 truncation constructs by APC-conjugated anti-Flag or isotype control antibodies. (C) Example of flow cytometry gating strategy used to quantify cells expressing GFP following infection by GFP-expressing RVPs. (D) Staining of K562 cells expressing the indicated alphavirus receptors. PE: *R*-phycoerythrin.

**Supplementary Figure 9.**
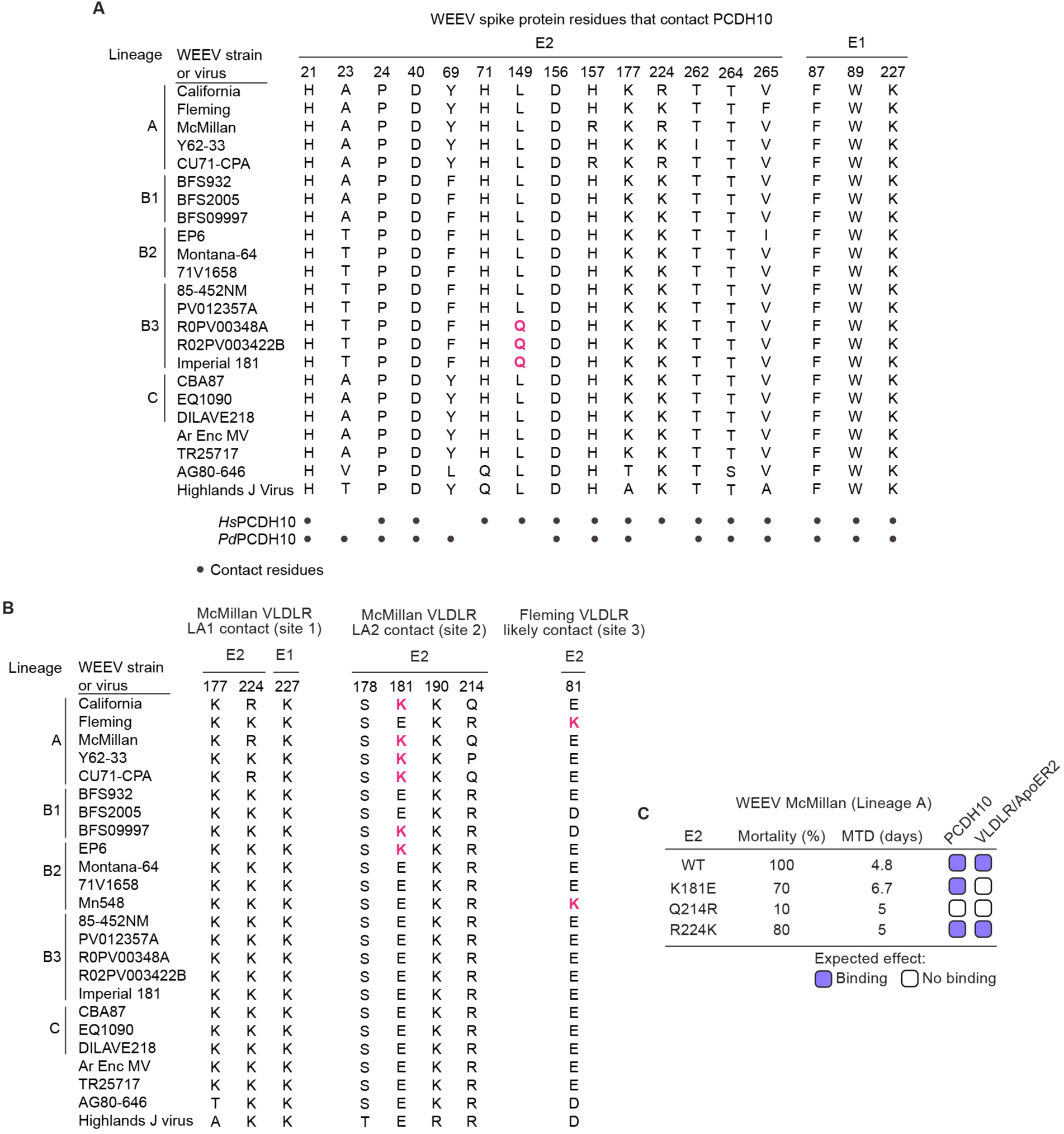
List of residues that contact receptors and summary of mouse virulence of McMillan mutants. (A) WEEV spike protein residues that contact *Hs*PCDH10 or *Pd*PCDH10 (< 4.0 Å). A key E2 polymorphic residue (Q149) is colored pink. (B) WEEV spike protein residues that contact VLDLR. Key polymorphic residues (E2 K181 and E2 K81) are colored in pink. (C) Effects of WEEV E2 McMillan mutations on mortality and mean time-to-death (MTD) when five-week-old female CD1 mice were inoculated 10^3^ PFU subcutaneously in the left thigh, summarized from Mossel et al.^38^ The expected effects of E2 mutations on PCDH10 or VLDLR/ApoER2 are based on the results of functional assays with McMillan RVPs shown in Figure 3G.

**Supplementary Figure 10.**
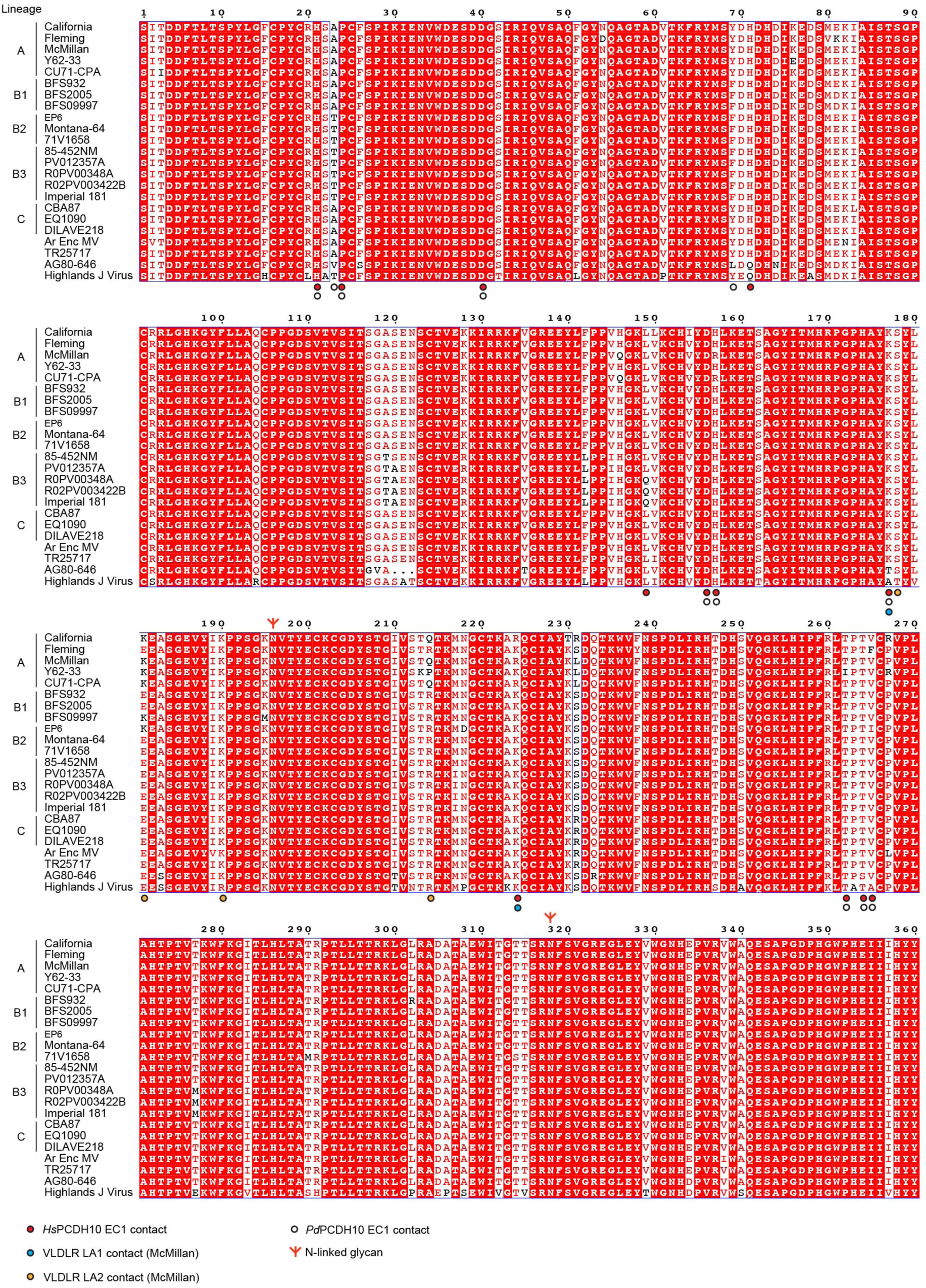
Sequence alignments of different WEEV E2 glycoproteins. Sequence alignments of the E2 glycoproteins from different WEEV strains. The strain information and accession numbers and are as follows: WEEV California (GenBank: KJ554965.1), WEEV Fleming (GenBank: MN477208.1), WEEV McMillan (GenBank: DQ393792.1), WEEV Y62-33 (GenBank: KT844544.1), WEEV CU71-CPA (GenBank: KT844545.1), WEEV BFS932 (GenBank: KJ554966.1), WEEV BFS2005 (GenBank: GQ287644.1), WEEV BFS09997 (GenBank: KJ554974.1), WEEV EP6 (GenBank: KJ554967.1), WEEV Montana-64 (GenBank: GQ287643.1), WEEV 71V1659 (GenBank: NP_640331.1), WEEV 85-452NM (GenBank: GQ287647.1), WEEV PV012357A (GenBank: KJ554987.1), WEEV R0PV00348A (GenBank: KJ554991.1), WEEV R02PV003422B (GenBank: KJ554990.1), WWEEV Imperial 181 (GenBank: GQ287641.1), WEEV CBA87 (GenBank: DQ432026.1), WEEV EQ1090 (GenBank: PP544260.1), WEEV DIAVE218 (GenBank: PP620644.1), WEEV Ar Enc MV (GenBank: KT844542), WEEV TR25717 (GenBank: KT844541), WEEV AG80-646 (GenBank: NC_075015), Highlands J virus 585-01 (GenBank: NC_012561.1). The Red background denotes residues that are completely conserved in all sequences. Boxed residues highlight positions where a single majority residue or multiple chemically similar residues are found. Such residues are highlighted in red. Receptor contact residues and N-linked glycan sites are indicated as shown in the legend. The panels were generated using ESPript 3.0.^54^

**Supplementary Figure 11.**
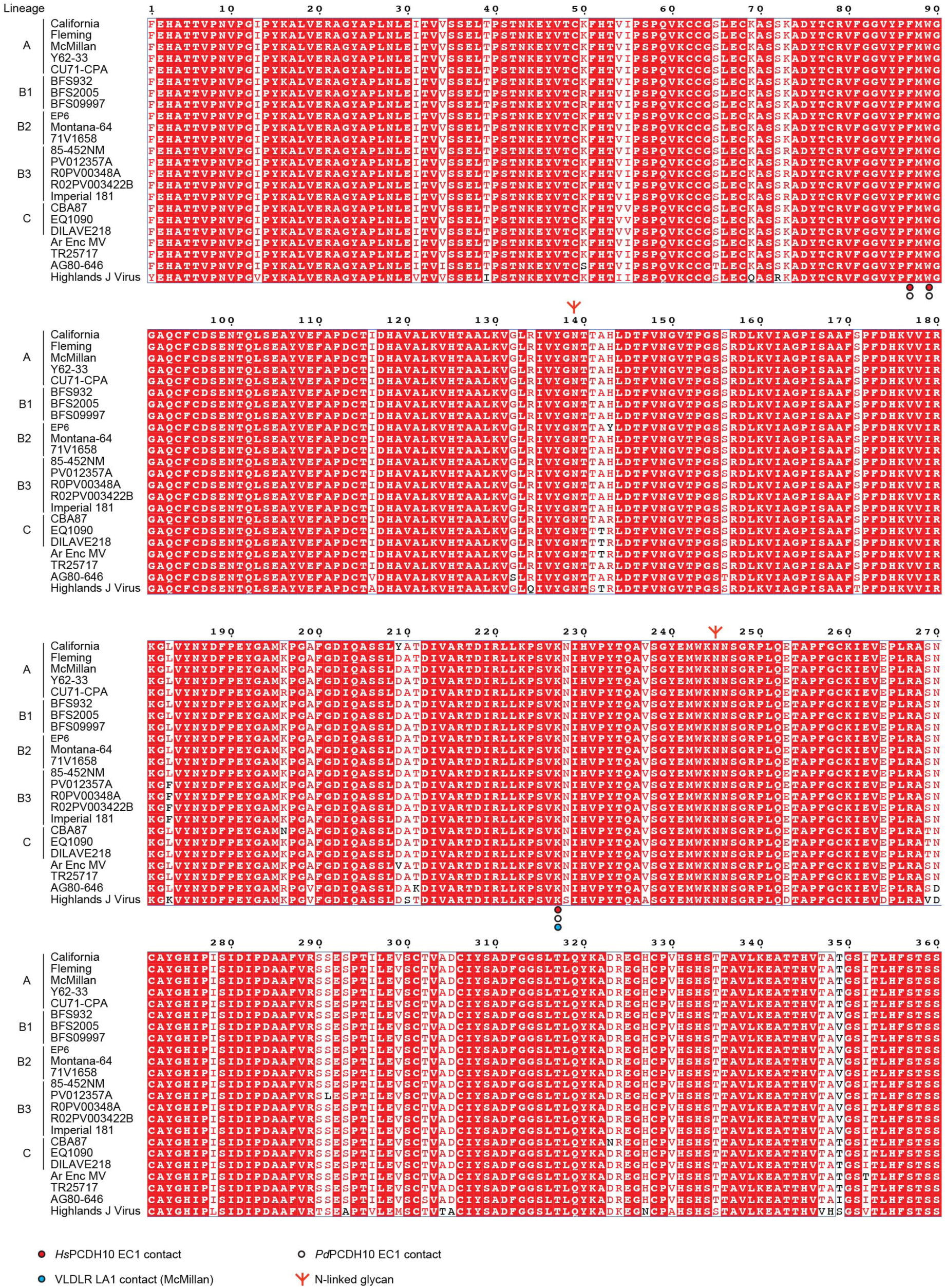
Sequence alignments of different WEEV E1 glycoproteins. Sequence alignments of the E1 glycoproteins from different WEEV strains. The strain information and accession numbers and are as follows: WEEV California (GenBank: KJ554965.1), WEEV Fleming (GenBank: MN477208.1), WEEV McMillan (GenBank: DQ393792.1), WEEV Y62-33 (GenBank: KT844544.1), WEEV CU71-CPA (GenBank: KT844545.1), WEEV BFS932 (GenBank: KJ554966.1), WEEV BFS2005 (GenBank: GQ287644.1), WEEV BFS09997 (GenBank: KJ554974.1), WEEV EP6 (GenBank: KJ554967.1), WEEV Montana-64 (GenBank: GQ287643.1), WEEV 71V1659 (GenBank: NP_640331.1), WEEV 85-452NM (GenBank: GQ287647.1), WEEV PV012357A (GenBank: KJ554987.1), WEEV R0PV00348A (GenBank: KJ554991.1), WEEV R02PV003422B (GenBank: KJ554990.1), WWEEV Imperial 181 (GenBank: GQ287641.1), WEEV CBA87 (GenBank: DQ432026.1), WEEV EQ1090 (GenBank: PP544260.1), WEEV DIAVE218 (GenBank: PP620644.1), WEEV Ar Enc MV (GenBank: KT844542), WEEV TR25717 (GenBank: KT844541), WEEV AG80-646 (GenBank: NC_075015), Highlands J virus virus 585-01 (GenBank: NC_012561.1). Red background highlights residues completely conserved in all sequences aligned. Boxed residues highlight positions where a single majority residue or multiple chemically similar residues could be identified. Such residues are highlighted in red. Receptor contact residues and N-linked glycan sites are indicated as shown in the legend. The panels were generated using ESPript 3.0.^54^

**Supplementary Figure 12.**
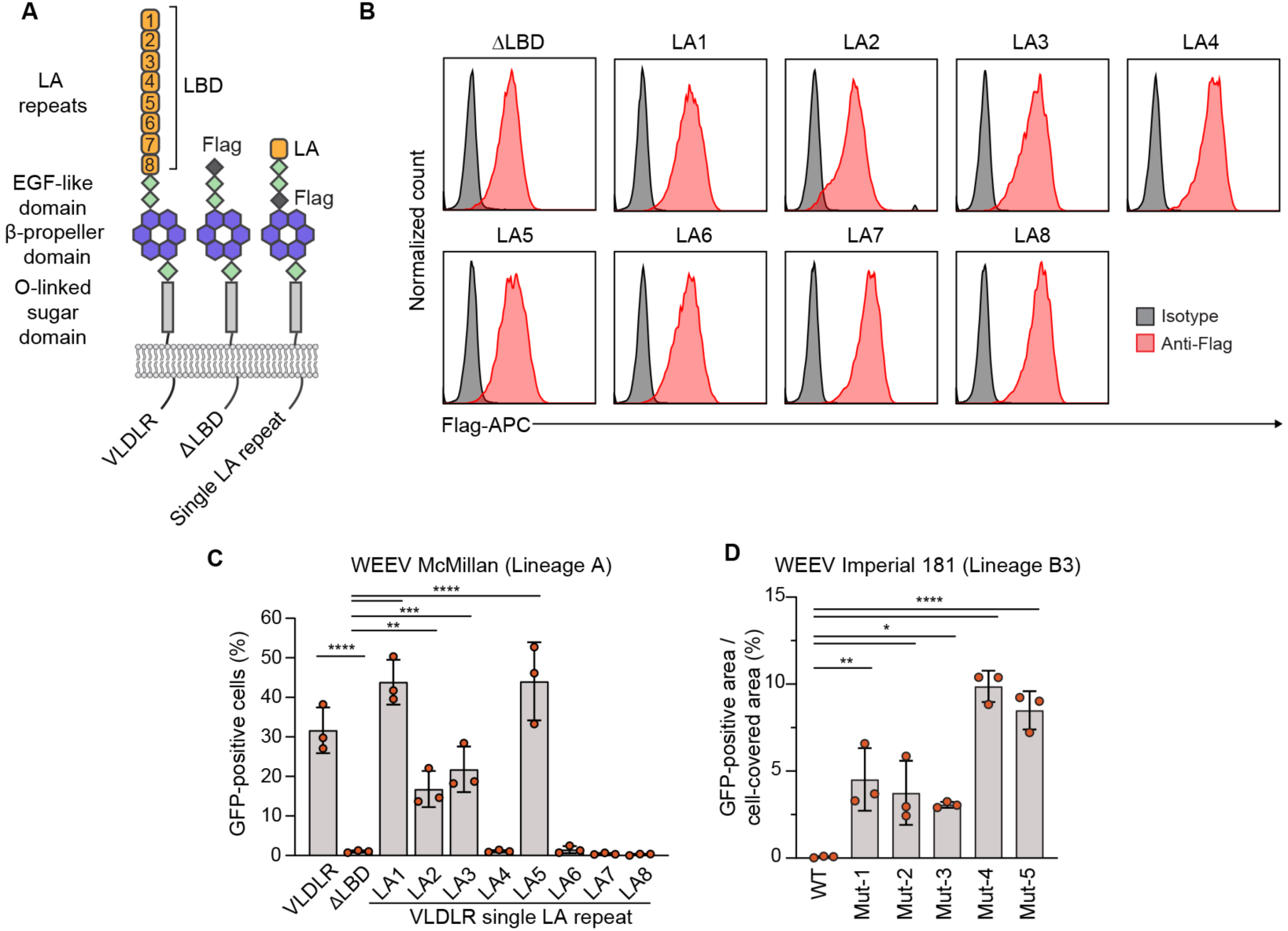
WEEV McMillan LA repeat preferences and infectivity results with WEEV Imperial 181 RVP mutants. (A) Schematic diagrams of wild-type VLDLR and single LA repeat constructs. A ΔLBD construct in which the entire LBD is replaced by an N-terminal Flag tag is used as a negative control in experiments. (B) Staining of K562 cell surface VLDLR or Flag-tagged VLDLR truncation constructs by APC-conjugated anti-Flag or isotype control antibodies. (C) Infection of K562 cells stably expressing VLDLR, ΔLBD-Flag, or single LA repeat constructs by WEEV McMillan RVPs. Infection was quantified by flow cytometry. (D) Primary murine cortical neurons were infected with GFP-expressing wild-type or mutant WEEV Imperial 181 RVPs. Infection was monitored using a live cell imaging system (see Methods for additional information). Data are mean ± s.d. from three experiments performed in triplicates (n=3) (C, D). One-way ANOVA with Dunnett’s multiple comparisons test, *\*\*\*\*P<*0.0001; \*\*\**P*<0.001; *\*\*P<*0.01 (C, D).

**Supplementary Figure 13.**
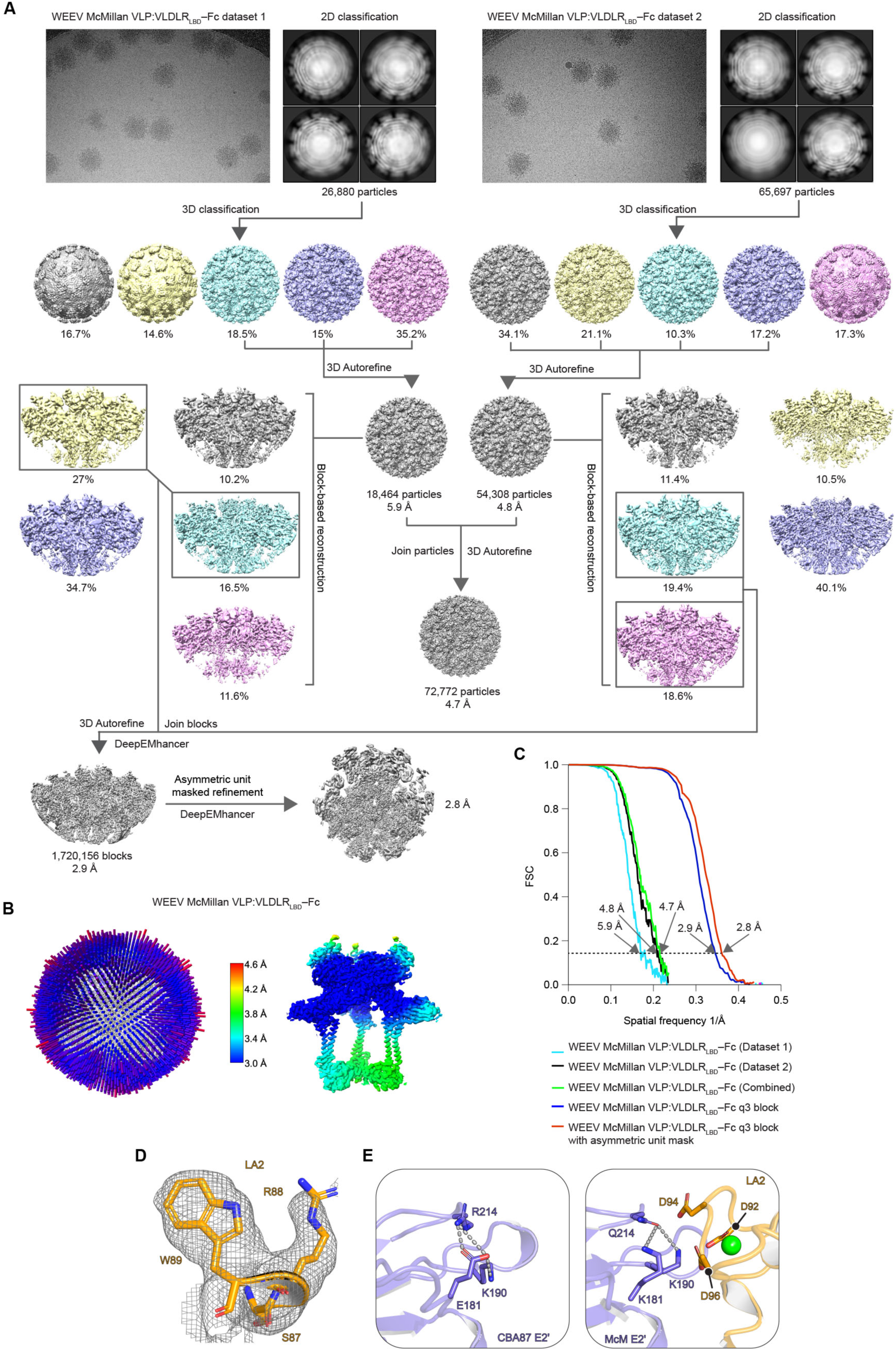
Cryo-EM reconstruction of WEEV McMillan VLP in complex with VLDLR_LBD_–Fc. (A) Workflow used for cryo-EM data processing of WEEV VLPs bound to VLDLR_LBD_–Fc. (B) 3D representation of particles angular distribution and local resolution maps of WEEV McMillan VLP in complex with VLDLR_LBD_–Fc. (C) Fourier shell correlation curves for WEEV McMillan VLP in complex with VLDLR_LBD_–Fc. The threshold used to estimate the resolution is 0.143. See Methods for additional details. (D) Fitting of VLDLR LA2 into the LA repeat density of site 2. An elongated side chain density that could only be explained by an arginine (present in LA2) but not a serine (present in LA3) allowed for unambiguous identification of LA2 as the bound LA repeat at this site. (E) Comparison of WEEV McMillan E2 and CBA87 E2 near the key polymorphic E2 residues at positions 181 and 214. E2 E181 in CBA87 forms a salt bridge with E2 R214 (left panel). The Q214R substitution in McMillan E2 may locally destabilize E2 by positioning three basic residues (K181, K190, and R214) in close proximity (right panel).

**Supplementary Figure 14.**
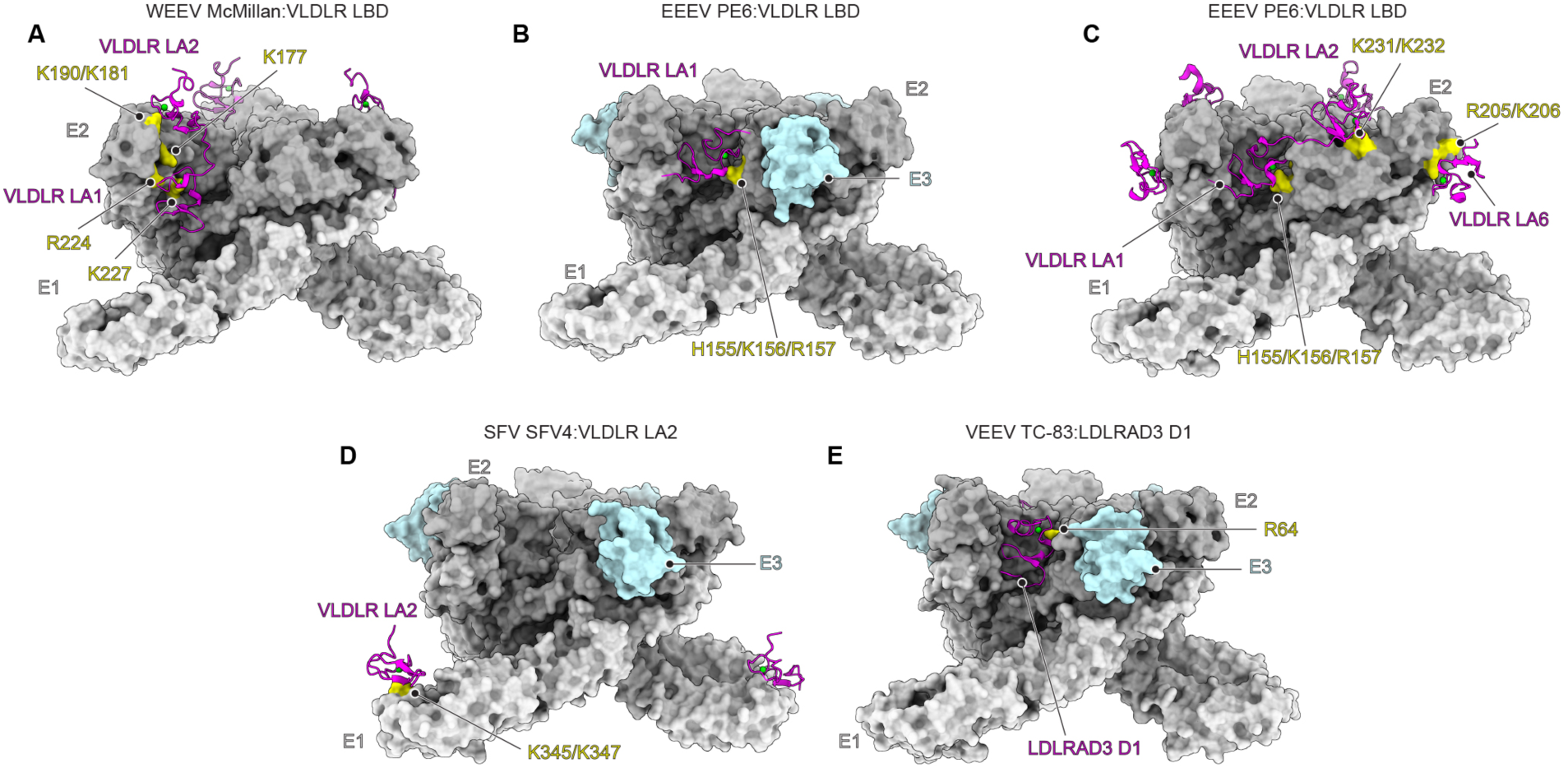
Comparisons of alphavirus E2 and E1 bound to LA repeats. (A–E) Alphavirus spike proteins are shown in a surface-rendered representation. The bound receptors are shown in ribbons. The LA repeat contact residues on WEEV McMillan, EEEV PE6 with retained E3 (B, PDB ID: 8UA4),^19^ EEEV PE6 (C, PDB ID: 8UFB),^20^ SFV SFV4 (D, PDB ID: 8UA8),^19^ and VEEV TC-83 (E, PDB ID: 7FFF)^17^ are colored in yellow. LA repeats are colored in magenta; E2–E1 glycoproteins are colored in different shade of gray; E3 is colored in light blue. LBD: ligand-binding domain. D1: domain 1.

**Supplementary Table 1.**
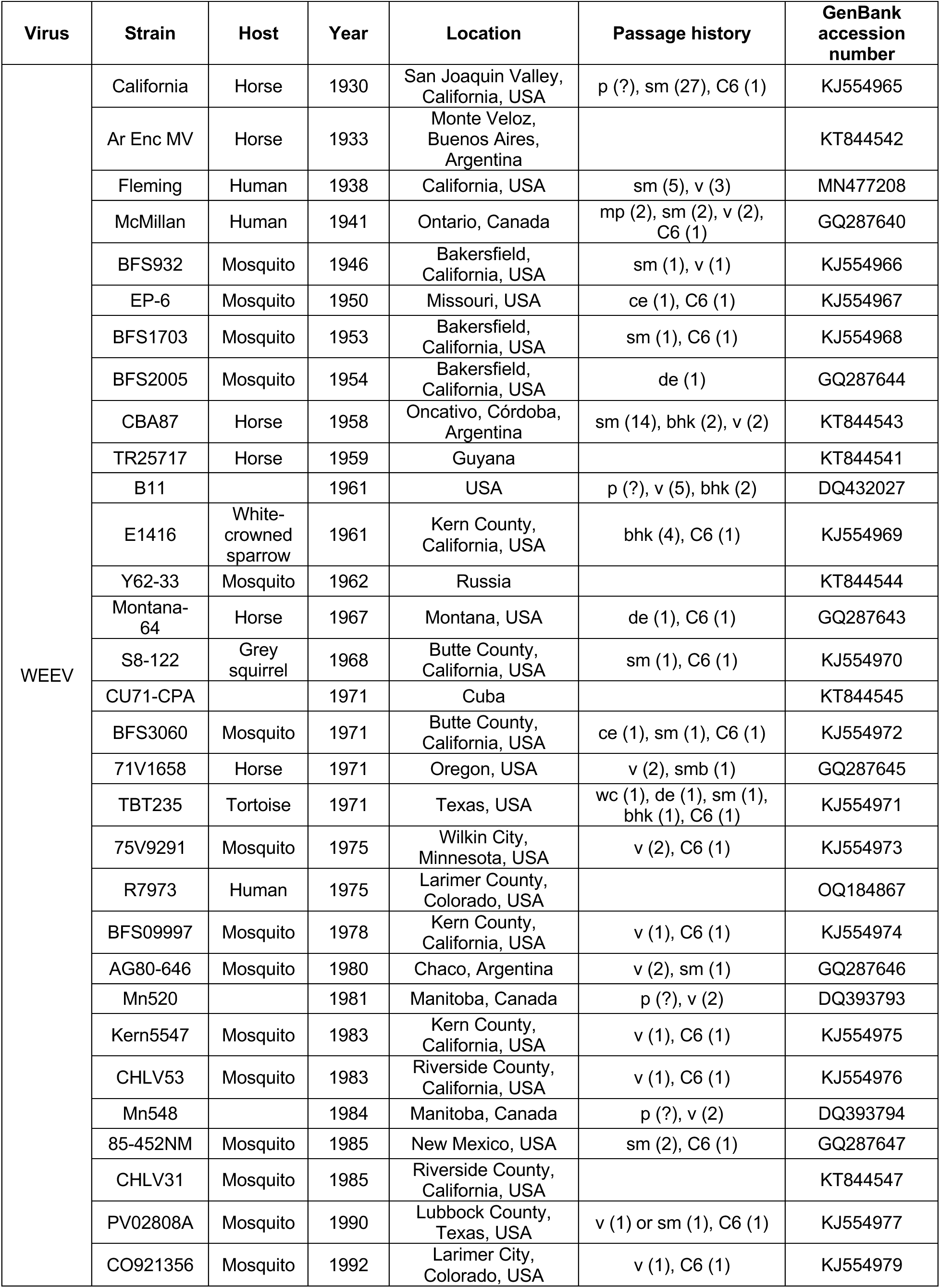
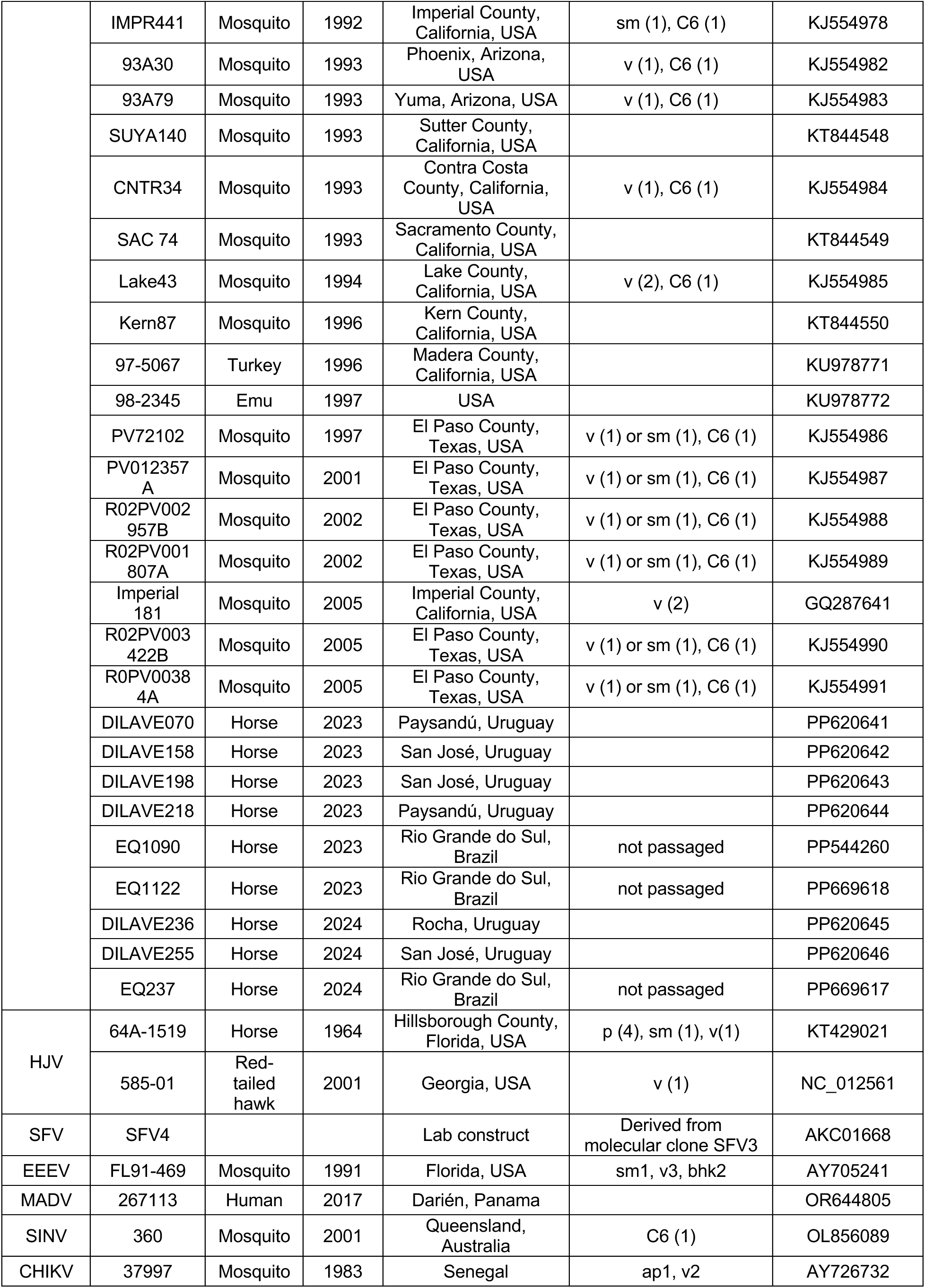
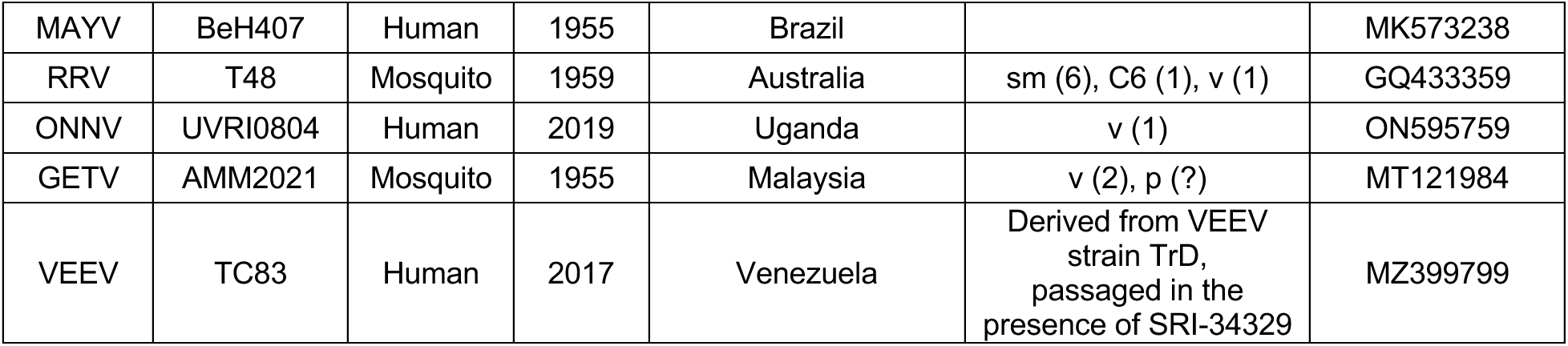
Strain information for viral sequences used in infectivity studies and phylogenetic analyses. Passage numbers are in parentheses. "?" indicates unknown passage number. Abbreviations: mp: mouse; sm: suckling mouse; smb: suckling mouse brain; v: Vero cells; bhk: baby hamster kidney cells; wc: wet chicks; de: duck embryonic fibroblasts; ce: chick embryonic fibroblasts; C6: C6/36 (*Aedes albopictus*) cells; ehe: embryonated hen’s eggs; gp: guinea pig; p: unknown media.

**Supplementary Table 2.**
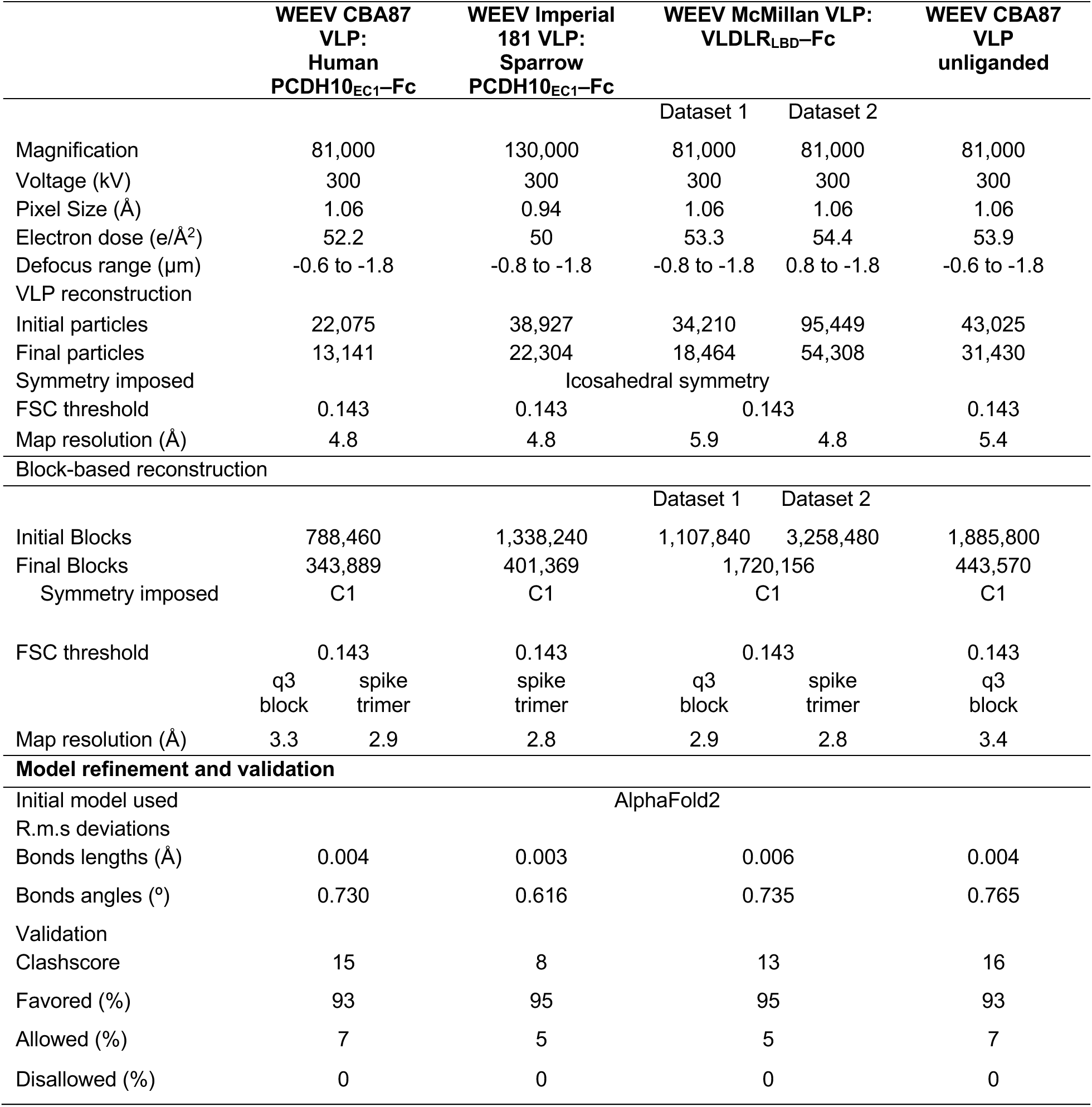
Cryo-EM data collection and validation statistics.

## Methods

### Cell lines and viruses

All cell lines used in this study are listed in the key resources table. HEK 293T (human kidney epithelial, ATCC CRL-11268), Vero E6 (*Cercopithecus aethiops* kidney epithelial, ATCC CRL-1586) were maintained in Dulbecco’s modified Eagle’s medium (DMEM, Gibco) supplemented with 10% (v/v) fetal bovine serum (FBS) and 25 mM HEPES (Thermo Fisher Scientific). K562 (human chronic myelogenous leukemia, ATCC CCL-243) cells were maintained in RPMI1640 (Thermo Fisher Scientific) supplemented with 10% (v/v) FBS, 25 mM HEPES, and 1% (v/v) penicillin-streptomycin. Expi293F cells (Thermo Fisher Scientific Cat#: A14527) were maintained in Expi293 Expression Medium (Thermo Fisher Scientific). Cell lines were not authenticated. Absence of mycoplasma is confirmed through routine mycoplasma test using e-Myco PCR detection kit (Bulldog Bio Cat#: 25234).

### Production and purification of virus-like particles

The genes encoding the structural polyprotein (capsid-E3-E2-(6K/TF)-E1) of WEEV strains CBA87 (GenBank: DQ432026.1), McMillan (GenBank: DQ393792.1), and Imperial 181 (GenBank: GQ287641.1) with capsid K67N mutation used to improve the yield of VLP production^30^ were separately cloned into the pVRC vector. We also produced EEEV VLPs using a previously described vector that encodes the structural polyprotein of EEEV strain PE6 (GenBank: AAU95735.1) with capsid mutation K67N.^30^ Expi293F^TM^ (Thermo Fisher Scientific) cells were cultured at 37°C with 5% CO_2_ in the Expi293^TM^ expression medium. We transfected Expi293F^TM^ cells with pVRC vectors encoding WEEV structural polyproteins using ExpiFectamine^TM^ 293 Transfection Kit (Thermo Fisher Scientific) according to the manufacturer’s instructions. We collected culture supernatant 5 d post-transfection and pelleted cells by centrifugation at 3,000 g for 20 min. The clarified supernatant was ultracentrifuged with a sucrose cushion consisting of 5 ml of 35% (w/v) sucrose and 5 ml of 70% (w/v) sucrose at 110,000 x *g* for 5 h in a Beckman SW28Ti rotor at 4°C. The VLPs were pooled from the interface of the 35% (w/v) and 70% (w/v) sucrose cushions, then buffer exchanged to lower the sucrose concentration to less than 20% (v/v) at a volume of 1 ml using a 100-kDa Amicon filter (Sigma). The VLPs were loaded onto a 20%–70% (v/v) continuous sucrose density gradient and centrifuged for 1.5 h in a Beckman SW41 rotor at 210,000 x *g* at 4°C. The VLP band was collected and exchanged into Dulbecco’s Phosphate Buffered Saline (DPBS, Thermo Fisher Scientific Cat#: 14190-144) using a 100-kDa Amicon filter. We confirmed integrity and purity of VLPs using SDS-PAGE (Figures S2A and S7A).

### Expression and purification of recombinant proteins

The genes encoding the EC1 domain of human PCDH10 (residues 19-122, GenBank NP_116586.1),^10^ the human VLDLR LBD (residues 31–355, GenBank NP_003374.3),^39^ or the human MXRA8 ectodomain (residues 20–337, GenBank NP_001269511.1)^39^ with the human IgG1 Fc region at their C terminal were separately cloned into the pVRC vector. We subcloned the EC1 domain of *P. domesticus* PCDH10 (residues 19-122, GenBank XP_064272571.1) into the same pVRC vector, provided by A. Schmidt.^66^ To purify PCDH10_EC1_–Fc and MXRA8_ect_–Fc, we transfected Expi293F^TM^ cells with plasmids encoding the fusion proteins using ExpiFectamine^TM^ 293 Transfection Kit (Thermo Fisher Scientific Cat#: A14525) according to the manufacturer’s instructions. Supernatants were collected 5 d post-transfection, centrifuged at 4,000 g for 30 min and purified with MabSelect^TM^ PrismA protein A affinity resin (Cytiva Cat#: 17549801) using the manufacturer’s protocol. The proteins were further purified by size-exclusion chromatography on a Superdex 200 increase 10/300 column (Cytiva). Proteins were stored in Tris Buffered Saline (TBS) (20 mM Tris, 150 mM NaCl in water, pH 7.5).

To purify VLDLR_LBD_–Fc, we co-transfected Expi293F cells with the pVRC vector encoding VLDLR_LBD_–Fc and a pCAGGS vector encoding the chaperon receptor-associated protein (RAP, residues 1-353, GenBank NP_002328). Supernatants were collected 5 d post-transfection and purified with MabSelect^TM^ PrismA protein A affinity resin. We separated the VLDLR_LBD_–Fc from RAP on the column by washing the column with 300 volumes of 10 mM EDTA in TBS overnight. Then VLDLR_LBD_–Fc was refolded on the column by washing with 100 column volumes of 2 mM CaCl_2_ in TBS and eluted using the manufacturer’s protocol. The proteins were concentrated and further purified by size-exclusion chromatography on a Superdex 200 increase 10/300 GL column. RAP was stored in TBS and VLDLR_LBD_–Fc was stored in TBS containing 2 mM CaCl_2_.

The genes encoding the EC1 domain of human PCDH10 (residues 19-122, GenBank NP_116586.1) were cloned into a pVRC vector with a C-terminal twin-strep tag for purification. We transfected Expi293F^TM^ cells with plasmid using ExpiFectamine^TM^ 293 Transfection Kit according to the manufacturer’s instructions. After centrifugation at 4,000 g for 30 min, supernatants were incubated with Strep-Tactin^®^ XT Sepharose resin (IBA Lifesciences) for 1 h at 4 °C. The beads were then washed using TBS, and the bound proteins were eluted using 50 mM biotin in TBS. The proteins were further purified by size-exclusion chromatography on a Superdex 200 increase 10/300 column equilibrated with TBS.

### Biolayer interferometry binding assays

We performed biolayer interferometry experiments with an Octet RED96e (Sartorius) and analyzed data using ForteBio Data Analysis HT version 12.0.1.55 software. The anti-WEEV E protein monoclonal antibody SKW11 (a gift from M. Sutton and M. Roederer)^56^ was loaded onto Anti-Human IgG Fc Capture (AHC) Biosensors (Sartorius Cat#: 18-5063) at a concentration of 250 nM in kinetic buffer (TBS supplemented with 2 mM CaCl_2_, 0.1% (w/v) BSA and 0.01% Tween) for 600 s. After a baseline measurement in kinetic buffer for 60s, tips were dipped into wells containing 1 μM WEEV CBA87 VLPs for 1 h. The signal was then allowed to equilibrate for an another 1 h in kinetic buffer. The tips were washed with the kinetic buffer for 60 s to obtain a baseline reading, then the biosensors were dipped into wells containing the serial dilutions of human PCDH10_EC1_–twin-strep (concentration range of 10 μM to 312.5 nM) in kinetic buffer for 900 s. Finally, a 300 s dissociation in kinetic buffer was performed. Data analysis was performed using a standard 1:1 binding model.

### ELISA

Apparent affinities of *Hs*PCDH10 EC1–Fc and *Pd*PCDH10_EC1_–Fc for CBA87, McMillan and Imperial 181 VLPs were also separately determined by ELISA. 200 ng VLPs were immobilized on ELISA MaxiSorp plates (Thermo Scientific Cat#: 439454) overnight at 4 °C. The coated plates were blocked by PBS supplemented with 3% (w/v) BSA for 1 h at room temperature, then were washed five times with PBS. Serial dilutions of *Hs*PCDH10_EC1_–Fc or *Pd*PCDH10_EC1_–Fc proteins were prepared and added to plates and incubated for 1 h at room temperature. After incubation with Fc-fusion proteins, the plates were washed five times with PBS, and then 100 μl per well of horseradish-peroxidase-conjugated anti-Human IgG (Sigma Cat#: A0170) diluted at ratio of 1:20,000 in PBS supplemented with 3% BSA were added for incubation for 1 h at room temperature. The plates were washed five times with PBS, then incubated with 100 μl per well of 1-step TMB ELISA solution for 3 min at room temperature in the dark and the reactions were stopped by addition of 100 μl per well of 2 N sulfuric acid. Absorbance was read at an optical density of 450 nm using BioTek multi-mode reader. EEEV VLPs were used as negative controls in experiments.

### Truncation and mutagenesis, and stable cell line generation

cDNA encoding human PCDH10 (GenBank NM_032961.3), human VLDLR (GenBank NP_003374.3), human MXRA8 (GenBank NM_032348.3), human ApoER2 isoform 2 (GenBank NM_004631.5), *P. domesticus* PCDH10, *P. domesticus* MXRA8, *E. caballus* PCDH10 (GenBank XM_023636548.1), *T. sirtalis* PCDH10 (GenBank XM_014072689.1), *M. musculus* PCDH10 (GenBank: NP_001091640.1) were cloned into a lentiGuide-Puro (Addgene #52963) expression vector as previously described.^10^ Non-human sequences were codon optimized for expression in human cells.

For the single PCDH10 EC1 construct, residues 19-690 in the PCDH10 precursor protein were replaced with residues 19-122 and adding a Flag tag (DYKDDDDK) between S696 and G697. Point mutations of contact residues in EC1 were generated using site-directed mutagenesis on the background of the single PCDH10 EC1 construct. For PCDH10 stalk-Flag, residues 19-690 in the precursor protein sequence were removed and a Flag tag (DYKDDDDK) was added between S696 and G697. VLDLR truncation constructs were previously described.^19^ Wild-type and mutant PCDH10 and VLDLR constructs were cloned into a lentiGuide-Puro (Addgene #52963) expression vector.

To generate lentiviruses for stable transduction, lentiGuide-Puro plasmids containing the transgene were co-transfected with psPAX2 (Addgene #12260) and PMD2.G (Addgene #12259) at a ratio of 3:2:1 into HEK 293T cells using Lipofectamine 3000 (Thermo Fisher Scientific). Lentiviruses were harvested 2 d post-transfection and used to transduce K562 cells. Successfully transduced K562 cells were selected for using puromycin at 2 μg ml^−1^. Cell lines were confirmed to express the constructs of interest at the plasma membrane by cell surface antibody staining (Figure S8). Antibodies are listed in the key resources table.

### Reporter virus particle generation

RVPs were generated as previously described^39^ using a dual vector system: a modified pRR64 Ross River virus replicon^67^ provided by R. Kuhn (Purdue University) (the SP6 promoter is replaced with a CMV promoter, the E3-E2-6K/TF-E1 sequence is replaced with a turbo GFP reporter preceded by a porcine teschovirus-1 2A self-cleaving peptide), and a pCAGGS vector expressing heterologous WEEV E3-E2-6K/TF-E1 proteins. The two vector were co-transfected into HEK 293T cells using Lipofectamine 3000 (Thermo Fisher). 4-6 h post-transfection, we replaced media with Opti-MEM (Thermo Fisher) supplemented with 5% (v/v) FBS, 25 mM HEPES, and 5 mM sodium butyrate. We harvested supernatant 2 d post-transfection, centrifuged supernatant at 4,000 rpm for 5 min, filtered using a 0.45 µm filter, and froze aliquots at −80 °C for storage.

WEEV E3-E2-6K/TF-E1 coding sequences cloned into the pCAGGS vector include: strain 71V1658 (GenBank NC_003908.1), strain Fleming (GenBank MN477208.1), strain McMillan (GenBank GQ287640.1), strain Imperial 181 (GenBank GQ287641.1), strain CBA87 (GenBank KT844543.1), strain BFS09997 (GenBank KJ554974.1), strain EP6 (GenBank KJ554967.1), strain AG80-646 (GenBank NC_075015.1), strain TR25717 (GenBank KT844541.1), strain EQ1090 (GenBank PP544260.1), strain DILAVE218 (GenBank PP620644.1). For HJV, strain 585-01 (GenBank NC_012561.1) was used. Mutant E3-E2-6K/TF-E1 sequences were generated using site-directed mutagenesis.

Titration of RVPs for MOI calculation was performed on Vero E6 cells seeded in a 96-well plate using a serial tenfold dilution of the RVP stocks. 24 h post-infection, numbers of GFP-positive cells were counted and used to calculate RVP titer as infectious unit per milliliter (IU ml^-1^), assuming that at high dilution factors, each GFP-positive cell is infected with one RVP given that RVPs only infect cells for one cycle.

### Reporter virus particle entry assays

We incubated transduced K562 cells with RVPs. 24 h post-infection, cells were harvested, washed twice with PBS, and fixed in PBS containing 2% (v/v) formalin. GFP expression was measured by FACS using an iQue3 Screener PLUS (Intellicyt) with IntelliCyt ForeCyt Standard Edition version 8.1.7524 (Sartorius) software. An example of the flow cytometry gating scheme used to quantify GFP-expressing RVP infection is provided in Figure S8.

For assessment of efficiency of receptor recognition, RVPs were titrated on K562 cells expressing cognate receptors in a twofold dilution series, with multiplicity of infection (MOI) calculated based on titers measured on Vero E6 cells.

### Mouse cortical neuron culture and infection

Embryonic day 17 mouse cortical neurons were purchased (Thermo Fisher Scientific Cat#: A15586) and cultured according to the manufacturer’s protocol with modifications. In brief, neurons were thawed and plated on 96 well plates coated with Poly-D-Lysine at 4.5 µg cm^-2^ at 40,000 neurons per well, in neurobasal medium (Thermo Fisher Scientific Cat#: 21103) supplemented with 0.5 mM Glutamax (Thermo Fisher Scientific Cat#: 35050), 2% (v/v) B-27 (Thermo Fisher Scientific Cat#: 17504), and 10 µM Y-27632 (Stemcell Technologies Cat#: 72302). 24 h post-plating, a full medium change was performed to remove Y-27632. Half medium change was performed every third day until infection. Neurons were infected with WEEV Imperial 181 wild-type or mutant RVPs at an MOI of 2 in the presence or absence of a control IgG (316 µg ml^-1^), PCDH10_EC1_–Fc (216 µg ml^-1^), or RAP (100 µg ml^-1^). 8 h post-infection, neurons were scanned using the Incucyte S3 Live Cell Imaging system (Sartorius) with Incucyte S3 Software version 2023A Rev2 (Sartorius) using a 20× objective. GFP-positive neurons were scored as cells with a threshold signal greater than 3 green calibrated units (GCU) above background, using a Surface Fit background subtraction method. The neuronal cell body area in each image was obtained by analyzing phase-contrast images using the Incucyte S3 Software (version 2023B). To calculate the percentage of positive cells, at the time point of 8 h post-infection, the area of GFP signal above background was divided by the total area covered by neuronal cell bodies and was multiplied by 100. Relative infection was calculated as follows: Relative infection (%) in the presence of recombinant proteins = (Percentage of GFP-positive cells in the presence of recombinant proteins) / (Percentage of GFP-positive cells under mock treatment) × 100.

### Cryo-EM sample preparation and data collection

We first mixed 3 μl of WEEV VLPs (with structural polyprotein sequence derived from strain CBA87 [GenBank: DQ432026.1]) at a concentration of 3.0 mg ml^-1^ in DPBS (Thermo Fisher Scientific Cat#: 14190-144) with 3 μl of *Hs*PCDH10_EC1_–Fc protein at 3.0 mg ml^-1^, then immediately applied 3 μl of sample to glow-discharged Quantifoil grids (R 0.6/1 300 mesh, gold, EMS Cat#: Q350AR-06) and blotted once for 5 s after a wait time of 15 s in 100% humidity at 4 °C and plunged into liquid ethane using an FEI Vitrobot Mark IV (Thermo Fisher Scientific). WEEV Imperial 181 VLPs (with structural polyprotein sequence derived from strain Imperial 181 [GenBank: GQ287641.1]) were also incubated with *Pd*PCDH10_EC1_–Fc for 4 h, and samples were frozen on Quantifoil grids (R 2/2 300 mesh, gold, EMS Cat#: Q3100AR2). WEEV McMillan VLPs (with structural polyprotein sequence derived from strain McMillan [GenBank: DQ393792.1]) were also incubated with VLDLR_LBD_–Fc, and samples were immediately frozen on Quantifoil grids. We also applied 3 μl of WEEV CBA87 VLP sample onto glow-discharged Quantifoil grids and plunged into liquid ethane to prepare cryo-EM samples with same parameters as WEEV VLP:PCDH10_EC1_– Fc complex using an FEI Vitrobot Mark IV. The Cryo-EM data collections were performed using a 300 kV FEI Titan Krios microscope (Thermo Fisher Scientific) with K3 (Gatan) or Falcon 4 (Thermo Fisher Scientific) direct electron detector at the Harvard Cryo-Electron Microscopy Center. Automated single-particle data acquisition was performed with SerialEM or EPU, with a magnification of ×81,000 (K3) or ×130,000 (Falcon 4) in counting mode, which yielded a calibrated pixel size of 1.06 Å (K3) or 0.94 Å (Falcon 4) and the datasets were collected at defocus ranges of −0.6 to −1.8 μm or −0.8 to −1.8 μm.

### Cryo-EM data processing

Raw movie stacks were corrected for beam-induced motion using MotionCor2 (version 1.6.4).^60^ The parameters of the contrast transfer function (CTF) for all micrographs were estimated by CTFDIND-4.1 (version 4.1.14).^61^ For the WEEV CBA87 VLP in complex with *Hs*PCDH10_EC1_–Fc, a total 22,075 particles were auto-picked from 7,749 micrographs using crYOLO (version 1.8.2),^58^ and particles were extracted with binned two times (pixel size 2.12 Å) in RELION 3.1 (version 3.1.4).^57^ After several rounds of reference-free 2D classification, 19,527 particles were selected from good 2D classes and subjected to 3D classification with icosahedral symmetry, and these particles were classified into five classes using the 12 Å density map of WEEV VLP (EMD-5210)^68^ as the initial reference model. After 3D classification, a total of 13,141 particles were selected and subjected to further 3D auto-refinement with icosahedral symmetry, finally yielding 4.8 Å reconstruction of the WEEV CBA87 VLP:*Hs*PCDH10_EC1_–Fc complex. To improve the resolution of the map, we used the block-based reconstruction method^69^ centered on the q3 spikes. 788,460 blocks were extracted and subjected to 3D classifications without alignment. 343,889 particles were selected for initial refinement with C1 symmetry, which was followed by CTF refinement, and a final round of refinement that generated a 3.3 Å map of the q3 spikes. We then used masked refinement to obtain a 2.9 Å density map of the spike trimer, which includes three copies of E2– E1 heterodimers and three copies of the capsid. This map was processed using DeepEMhancer (version 20210511)^59^ for cryo-EM volume post-processing to generate a map for model building. The cryo-EM WEEV VLP alone data were processed with similar data processing procedure using RELION 3.1 (version 3.1.4),^57^ which generated a 5.4 Å map of the WEEV CBA87 VLP alone. Then we got a 3.4 Å map of the q3 spikes by block-based reconstruction method.^69^ This map was processed using DeepEMhancer (version 20210511)^59^ for cryo-EM volume post-processing to generate a map for model building. And the cryo-EM WEEV Imperial 181 VLP bound to *Pd*PCDH10_EC1_–Fc data were processed with similar data processing procedure using RELION 3.1 (version 3.1.4),^57^ which generated a 4.8 Å map of the WEEV Imperial 181 VLP bound to *Pd*PCDH10_EC1_–Fc. We then obtained a 2.8 Å map of the trimeric spikes bound to *Pd*PCDH10_EC1_–Fc using the block-based reconstruction method.^69^

For the WEEV McMillan VLP in complex with VLDLR_LBD_–Fc, we processed the first dataset with similar data processing steps and get a 5.9 Å map of the WEEV McMillan VLP:VLDLR_LBD_– Fc. To improve the quality of density map, we collected a second dataset. The second dataset was processed separately and generated a 4.8 Å map of the WEEV McMillan VLP:VLDLR_LBD_– Fc. We then combined VLPs from both datasets after the first round of refinement and subjected them to refinement. Finally, the resolution of the combined WEEV McMillan VLP:VLDLR_LBD_–Fc complex map was 4.7 Å. We performed block-based reconstruction of q3 spikes of both datasets independently. We combined good blocks in the 3D classification results of both datasets and 1,720,156 blocks were used to perform 3D auto-refinement, which generated a 2.9 Å map of the q3 spikes. To improve features for the LA repeat density, an asymmetric unit masked refinement was performed. The resolution of masked refinement map is 2.8 Å, and DeepEMhancer-post-processing yielded a density map for model building with details of the spike protein, capsid, and that were well-resolved. Maps were processed using DeepEMhancer (version 20210511)^59^ for cryo-EM volume post-processing.

### Model building

For the WEEV CBA87 VLP:*Hs*PCDH10_EC1_–Fc, we used coordinates of PCDH10 EC1 from the crystal structure of human PCDH10 EC1–EC4 (PDB:6VFQ)^14^ and of WEEV E2–E1 and capsid predicted by AlphaFold2^65^ as initial models. These models were docked into the DeepEMhancer post-processed cryo-EM density map using UCSF ChimeraX (version 1.6.1).^62^ The atomic model was then generated through iterative rounds of model building and adjustment in *Coot* (version 0.9.8.91)^63^ and refined using real space refinement in Phenix (version 1.21rc1-5127).^64^

The model of the unliganded WEEV CBA87 VLP was generated based on WEEV CBA87 VLP:*Hs*PCDH10_EC1_–Fc model that was docked into the DeepEMhancer post-processed cryo-EM density map of the unliganded VLP using UCSF ChimeraX. For the WEEV Imperial 181 VLP:*Pd*PCDH10_EC1_–Fc, the structure CBA87 VLP:*Hs*PCDH10_EC1_–Fc as an initial model was fitted into the DeepEMhancer post-processed cryo-EM map of the q3 spikes for the Imperial 181 complex. The initial coordinates of WEEV McMillan VLP:VLDLR_LBD_–Fc were generated based on WEEV CBA87 VLP:*Hs*PCDH10_EC1_–Fc model and VLDLR LA1–LA2 structure predicted by AlphaFold2.^65^ These models were docked into the cryo-EM density for the WEEV McMillan: VLDLR_LBD_-Fc complex using UCSF ChimeraX.

The atomic models for the unliganded WEEV CBA87 VLP, WEEV Imperial 181 VLP:*Pd*PCDH10_EC1_–Fc, WEEV McMillan VLP:VLDLR_LBD_–Fc were generated through iterative rounds of model building and adjustment with their associated maps in *Coot* (version 0.9.8.91)^63^ and refined using real space refinement in Phenix (version 1.21rc1-5127).^64^ All models were validated using RCSB PDB and refinement statistics are provided in Table S2. Figures of structural presenting were made by either UCSF ChimeraX (version 1.6.1)^62^ or PyMOL (version 3.0.2) (https://www.pymol.org/pymol). Software used in this project was curated by SBGrid.^70^

## QUANTIFICATION AND STATISTICAL ANALYSIS

### Statistical analysis

Data were deemed statistically significant when P values were <0.05 using version 10 of GraphPad Prism. Experiments were analyzed by one-way or two-way ANOVA with multiple comparison correction.

## Author contributions

X.F. produced VLPs, recombinant Fc-fusion proteins, performed biolayer interferometry and ELISA experiments, determined cryo-EM structures, and built and refined atomic models. W.L. generated cell lines, RVPs, and recombinant proteins, and performed infectivity studies and phylogenetic analysis. J.S.P. generated cell lines and RVPs and performed infectivity studies. J.O., L.V.T., and L.M.M., generated RVPs and performed infectivity studies. C.J.M., H.V., and T.K.B. produce recombinant Fc-fusion proteins. V.B. and P.Y. helped generate WEEV CBA87 VLPs. J.M.B., S.N., H.U., H.B., S.Y.C. and I.M.C. participated in experimental design and helped analyze and interpret data. W.M.dS. and S.C.W. performed ancestral sequence reconstruction. K.S.P. and J.A.P. analyzed and interpreted data. J.A. acquired funding and supervised the research. X.F., W.L., and J.A. wrote the original draft of the manuscript, and all authors participated in reviewing and editing the manuscript.

## Acknowledgements

Cryo-EM data were collected at the Harvard Cryo-Electron Microscopy Center for Structural Biology. We thank Harvard Cryo-EM Center staff members R. Walsh, S. Rawson, M. Mayer, C.Leistner, and R. Nair for assistance with cryo-EM data collection. J.A. is a recipient of Burroughs Wellcome Fund Investigators in the Pathogenesis of Infectious Disease Award. This work was also supported by a Vallee Scholar Award (J.A.), Smith Family Foundation Odyssey Award (J.A.), Charles E.W. Grinnell Trust Award (J.A.), NIH grant R01 AI182377 (J.A.), a G. Harold and Leila Y. Mathers Foundation Award (J.A.), NIH grant T32AI700245 (C.J.M., J.S.P., T.K.B., L.V.T. and J.O.), T32GM144273 (H.V.), T32AG000222-33 (H.B., S.Y.C.), NIH grant R24 AI120942 (S.C.W.), Jackson-Wijaya Fund (I.M.C.), Burroughs Wellcome Fund Investigators in the Pathogenesis of Infectious Disease Award (I.M.C.), Wellcome Trust grant 226075/Z/22/Z (W.M.dS), and NIH grant R01 MH125162 (H.U.).

